# Proteasomal surveillance of exposed cysteines

**DOI:** 10.64898/2026.09.29.755535

**Authors:** Elijah L. Mena, Ken C. Dong, Anthony Kairouz, Katherine A. Donovan, Mostafa A. Soliman, Hai-Tsang Huang, Bercin Yildiran, Eric S. Fischer, Richard I. Sherwood, Richard T. Timms, Andreas Martin, Stephen J. Elledge

**Affiliations:** Department of Genetics, Howard Hughes Medical Institute, Harvard Medical School, Boston, MA, USA; Division of Genetics, Department of Medicine, Brigham and Women’s Hospital, Boston, MA, USA; Department of Molecular and Cell Biology, Howard Hughes Medical Institute, University of California at Berkeley, Berkeley, CA, USA; California Institute for Quantitative Biosciences, University of California at Berkeley, Berkeley, CA, USA; Department of Cancer Biology, Dana-Farber Cancer Institute, Boston, MA, USA; Department of Biological Chemistry and Molecular Pharmacology, Harvard Medical School, Boston, MA, USA; Broad Institute of Harvard and MIT, Cambridge, MA, USA; Department of Medical Oncology, Harvard Medical School, Dana-Farber Cancer Institute, Boston, MA, USA; Cambridge Institute of Therapeutic Immunology and Infectious Disease, Department of Medicine, University of Cambridge, Cambridge, UK

**Keywords:** 26S proteasome, UPS, ubiquitin-independent degradation, cysteine, ornithine decarboxylase (ODC), FOS, zinc, proteostasis

## Abstract

Cysteine residues are critical for both the catalytic activity and structural stability of many proteins, yet their high chemical reactivity threatens proteome integrity. Whether cells have specialized mechanisms to recognize and eliminate proteins with aberrant or exposed cysteines is unknown. By systematically analyzing the stability of thousands of cysteine-containing peptides, here we identify two complementary surveillance mechanisms targeting proteins with exposed cysteine residues. First, a set of E3 ubiquitin ligases – including AMBRA1, TRPC4AP, FBXO3, and FBXO21 – detects double cysteines and targets proteins for ubiquitin-dependent proteasomal degradation. Second, the PSMD14/Rpn11 subunit of the 26S proteasome recognizes exposed cysteines in a ubiquitin-independent manner. Both pathways function in quality control to clear misfolded proteins exposing cysteine residues, including zinc finger-containing proteins. Mutational escape from the ubiquitin-independent pathway through dysregulation of ornithine decarboxylase or FOS levels is responsible for two human diseases, Bachmann-Bupp Syndrome and osteoblastoma, respectively. Our findings reveal conserved mechanisms for degrading proteins with exposed cysteines and uncover an alternative mode of substrate recognition by the 26S proteasome.

**Short Summary:** Cysteine exposure acts as a conserved degradation signal that triggers two modes of turnover by the 26S proteasome: ubiquitin-dependent and direct, ubiquitin-independent degradation. These findings demonstrate the importance of maintaining cysteine homeostasis and reveal an alternative mode of proteasomal degradation.

## INTRODUCTION

Maintaining proteome integrity requires the cell to efficiently recognize and eliminate misfolded or damaged proteins. This surveillance often focuses on recognizing hydrophobic residues, which are normally buried within folded proteins and when inappropriately exposed can lead to toxic protein aggregation. But exposed hydrophobic residues are not the only threat to protein homeostasis. In particular, cysteine residues are essential for enzymatic activity, metal coordination, and structural stabilization (fig. S1A) (*1*), but their high reactivity makes them vulnerable to oxidative damage. Indeed, cysteine oxidation has been implicated in pathologies ranging from diabetes to neurodegeneration and aging (*2*–*5*). Evolution has strongly selected against solvent-exposed cysteines, even more so than for hydrophobic residues (*6*). However, whether cells possess active surveillance mechanisms to detect and eliminate proteins with aberrant cysteine exposure remains unknown.

Most substrates destined for degradation are marked by ubiquitylation and subsequently cleared by the 26S proteasome (*7*). Nonetheless, a subset of proteins, such as FOS and ornithine decarboxylase (ODC), bypass the ubiquitin system and are degraded directly by the proteasome through less well-understood mechanisms (*8*–*12*). Although some ubiquitin-independent substrates rely on adaptor proteins or ubiquitin-like modifiers, including Midnolin and FAT10 (*13*–*15*), the full scope and molecular logic of ubiquitin-independent proteasomal degradation remains unknown.

Here, we identify specialized cellular systems that recognize and degrade proteins bearing exposed cysteines. Through high-throughput genetic screening, we identify (i) a cohort of E3 ubiquitin ligases that detect and ubiquitylate proteins harboring double cysteines, and (ii) a direct, ubiquitin-independent pathway in which the 26S proteasome engages unfolded protein segments with exposed cysteines through its PSMD14/Rpn11 subunit. Our results reveal conserved mechanisms that protect against the chemical liabilities of cysteine residues, and we define a noncanonical mode of substrate recognition by the proteasome.

## RESULTS

### A high-throughput protein stability screen identifies substrates of cysteine-dependent degradation

To systematically study cysteine-dependent degradation, we synthesized an oligonucleotide library encoding 34-amino acid peptides encompassing cysteine-containing sequences from human proteins, capturing most CXXC and CXC motifs in the proteome (fig. S1B). We prioritized these double cysteine motifs because they are typically highly conserved, commonly coordinate metal ions or mediate redox chemistry, and have been implicated in selective protein degradation (*6, 16*–*19*). Each cysteine-containing sequence was present in the expression library as three distinct tiles, offset in the original protein by 5 amino acids relative to one another. Then for each of 43,725 peptides representing wild-type sequences, we also synthesized a reference peptide in which a single cysteine residue was mutated to alanine. Peptides were fused to the C-terminus of GFP within the dual-color Global Protein Stability (GPS) reporter system (*20*) that also co-expresses DsRED as an internal control. Thus, protein stability could be assessed by measuring the GFP:DsRED ratio. We transduced HEK-293T cells, sorted them into six stability bins based on the GFP:DsRED ratio, and analyzed the composition of each bin by Illumina sequencing. From these data, we calculated a Protein Stability Index (PSI) score for each GFP-peptide fusion, ranging from 1 (maximally unstable) to 6 (maximally stable) based on the distribution of sequencing reads across the six stability bins (Fig. 1A). We performed two replicates of the screen, which were highly correlated (r = 0.97) (Fig. 1B).

**Fig 1.**
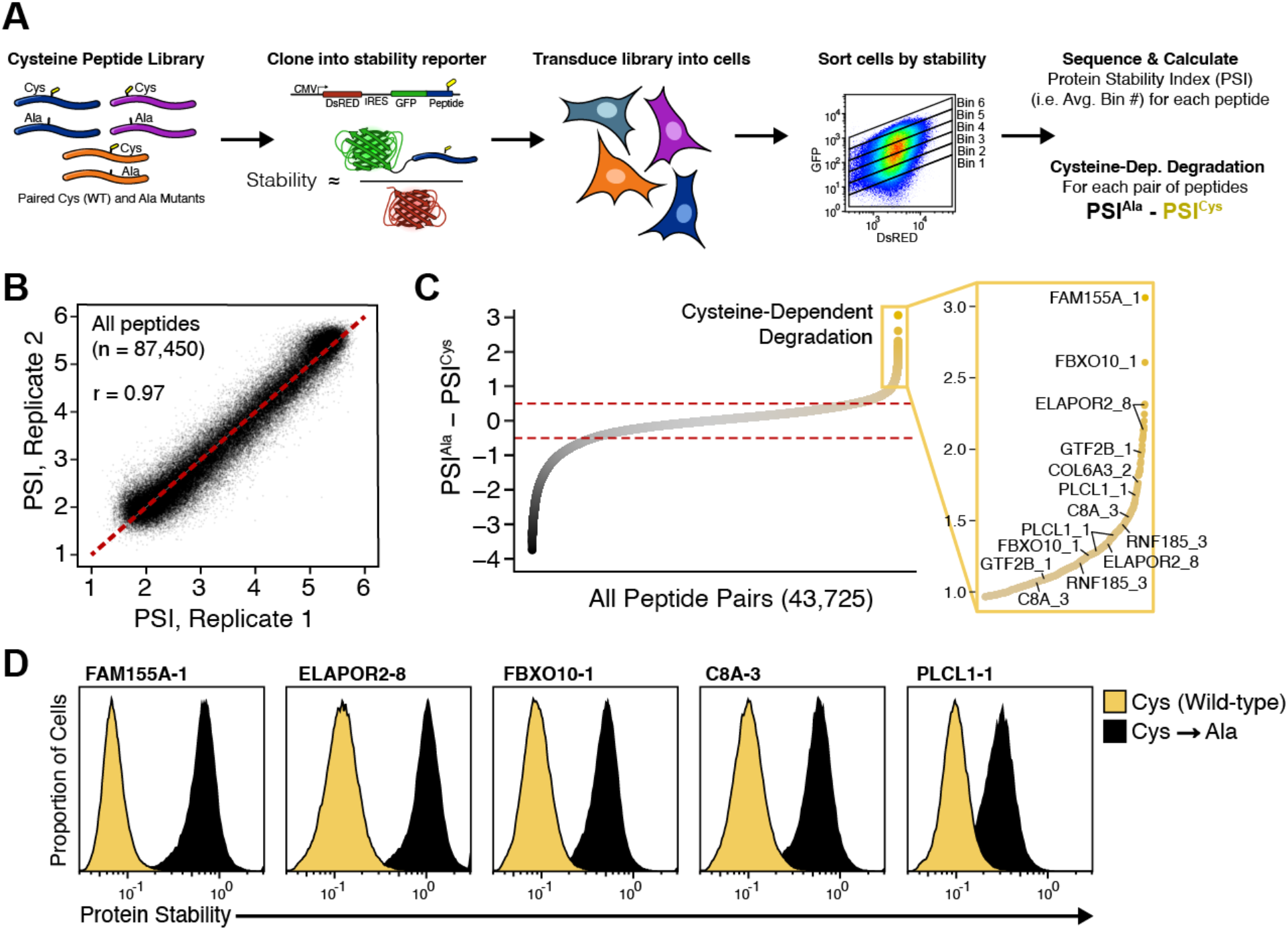
Specific cysteine residues trigger degradation. (**A**) Cysteine-containing 34-mer peptides derived from the human proteome were assessed for cysteine-dependent degradation using the Global Protein Stability (GPS) reporter system. (**B**) Protein stability index (PSI) values were highly correlated between biological replicates. (**C**) Cysteine-dependent degradation scores (PSI^Ala^ - PSI^Cys^) were calculated for all peptide pairs. Multiple overlapping peptides often consistently scored as top cysteine-dependent hits. (**D**) Selected cysteine-dependent degrons were validated using individual GPS reporters and flow cytometry. Substitution of the critical cysteine with alanine abolished degradation.

We calculated a cysteine-dependence score for each peptide pair (PSI^Ala^ - PSI^Cys^). Cysteine-to-alanine substitutions dramatically altered the stability of many sequences: 5,226 peptides (12%) were stabilized by ≥0.5 PSI units, indicating cysteine-dependent degradation (Fig. 1C). Leading candidates exhibiting cysteine-dependent degradation were concordantly identified across both replicate experiments (fig. S1C), often in multiple overlapping tiles (Fig. 1C, fig. S1D–I). Furthermore, individual validation assays using GPS confirmed that top-ranked substrates such as FAM155A-1 and ELAPOR2-8 were highly unstable, and that mutation of a single cysteine to alanine was sufficient to abolish this effect (Fig. 1D). Thus, cysteine exposure can strongly influence protein stability.

To define the cysteine-containing degrons at high resolution, we employed site-saturating mutagenesis. We selected 66 of the top cysteine-dependent peptides for mutagenesis, including the top hits as well as lower-scoring hits in which all three overlapping tiles scored. We then replaced each residue with every other amino acid and measured the impact on stability in a pooled screen. (Fig. 2A). This revealed two major classes of degrons. The instability of thirty-five peptides required two closely spaced cysteines, often with additional cysteines or other residues contributing (Fig. 2B, fig. S2A-H). In contrast, the instability of thirty-one substrates depended on only a single cysteine residue (Fig. 3A-C, fig. S2I-P). Therefore, there are two major classes of cysteine degrons: “double” cysteine degrons that require two cysteines and “single” cysteine degrons that contain one critical cysteine.

**Fig 2.**
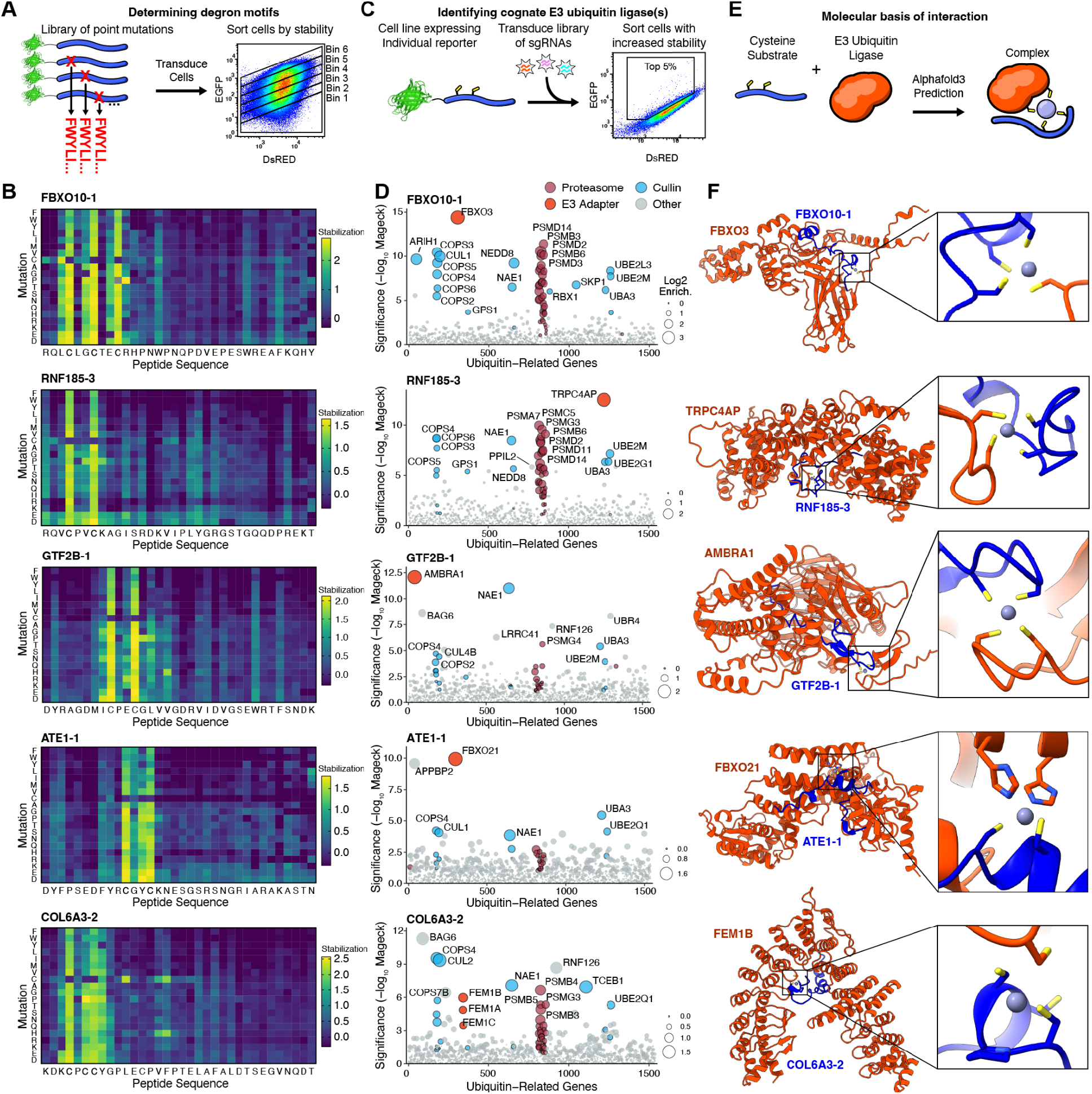
A family of Cullin-RING ligases target double cysteine motifs. (**A**) Site-saturating mutagenesis was performed on selected cysteine-dependent substrates. Mutant libraries were stably expressed in cells, followed by pooled stability sorting and high-throughput sequencing. (**B**) Heatmaps show mutational effects on representative double cysteine degrons. The wild-type peptide sequence is shown on the X-axis, with all single-residue substitutions on the Y-axis. Mutations that stabilize the degron (increase PSI) are shown in yellow. (**C**) CRISPR knockout screens were performed using a ubiquitin-focused sgRNA library to identify genes required for degradation of double cysteine substrates. Cells expressing stabilized peptides were sorted and sequenced for sgRNA enrichment. (**D**) Results from five CRISPR screens. Each dot represents a gene, dot size indicates sgRNA enrichment, and statistical significance (−log_10_[MAGeCK positive score]) is shown on the Y-axis. The proteasome, Cullin-related genes, and the scored E3 ligase are colored. (**E**) AlphaFold 3 modeling was used to predict interactions between paired-cysteine substrates and their cognate E3 ligases. (**F**) AlphaFold 3 results, with insets showing the cysteine-E3 interface.

**Fig 3.**
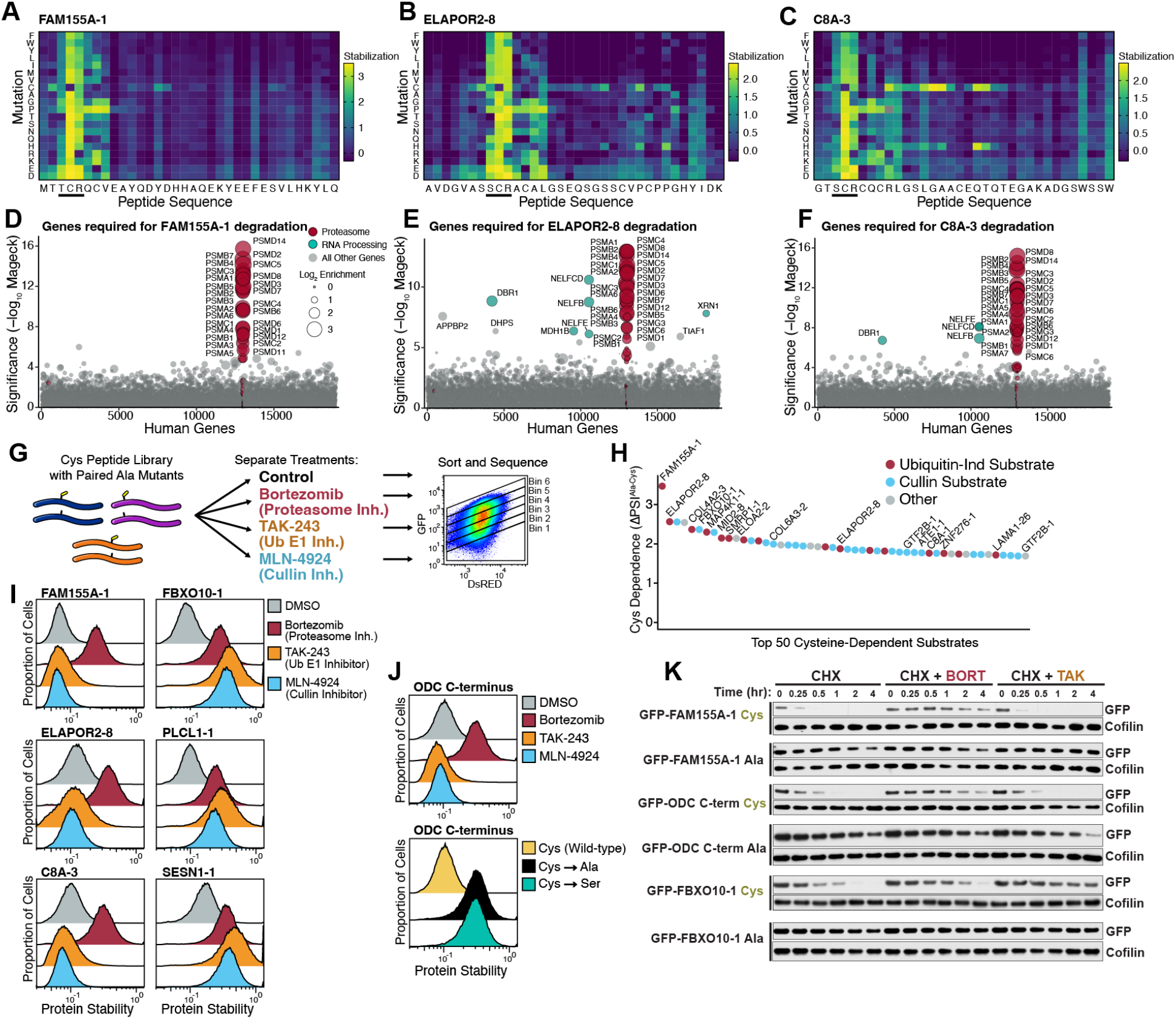
Single cysteines trigger proteasomal degradation through a ubiquitin-independent pathway. (**A**-**C**) Site-saturating mutagenesis reveals that degradation of three top-scoring substrates depends on a single cysteine residue (**D**-**F**) Genome-wide CRISPR-Cas9 screens performed on single-cysteine substrates identified the 26S proteasome as essential for degradation, with few other genes scoring. (**G**) The full cysteine-containing peptide library was re-screened for stability in the presence of ubiquitin-proteasome system inhibitors. (**H**) Most top-scoring cysteine-dependent substrates fall into two classes: ubiquitin-independent or Cullin-dependent. (**I**) Individual GPS reporter assays confirm that FAM155A-1, ELAPOR2-8, and C8A-3 are degraded in a bortezomib-sensitive but TAK-243-insensitive manner, indicating ubiquitin-independent proteasomal degradation. FBXO10-1, PLCL1-1, and SESN1-1 are double cysteine degrons and serve as positive controls. DMSO conditions were from the same experiment as Fig. 1D. (**J**) The C-terminal 37 amino acids of ODC confer ubiquitin-independent, cysteine-dependent degradation when appended to GFP. Residue Cys441 was mutated to alanine or serine. (**K**) Cycloheximide chase experiments were used to monitor protein stability over time in the presence of bortezomib or TAK-243.

To identify E3 ubiquitin ligases targeting these degrons, we performed 16 CRISPR knockout screens, each with a different cell line expressing an individual substrate subject to cysteine-dependent degradation. Following gene disruption using a ubiquitin-focused sgRNA library, we used FACS to isolate mutant cells in which these substrates were stabilized, and identified enriched sgRNAs by sequencing (Fig. 2C). For each of the 11 double cysteine degrons that we screened, we readily identified a candidate E3 ligase (Fig. 2D, fig. S3B), whereas none of the 5 single cysteine degrons that we screened were successful in identifying a putative E3 ligase (fig. S4). We first consider the double cysteine degrons in more detail.

### Double cysteine degrons are recognized by a family of Cullin-RING ligases

Across the 11 double cysteine substrates that we screened, we collectively identified seven distinct E3 ligases (Fig. 2D, fig. S3B): FBXO21, FBXO3, AMBRA1, TRPC4AP, and the FEM1A/B/C family. Cullin scaffold components, E2 enzymes, neddylation enzymes, and proteasomal genes were also consistently enriched, validating the robustness of our screening approach at identifying critical components of the degradation machinery. In each case, inhibiting Cullin-RING ligases (with MLN4924) or the 26S proteasome (with Bortezomib) conferred stabilization (fig. S3C), as did knockout of cognate E3 ligases identified by the screen (fig. S3D).

To understand how the E3 ligases recognize double cysteine degrons, we modeled substrate-E3 complexes using AlphaFold 3 (Fig. 2E). For each substrate, the models suggested that zinc acts as a molecular glue, bridging cysteines from the substrate to cysteines on the E3 ligase (Fig. 2F). Consistent with this mechanism, the zinc chelator TPEN stabilized several substrates (fig. S3C). Finally, we identified the E3 ligase APPBP2 as an E3 ligase that targets additional cysteine-containing substrates using an analogous zinc-mediated interaction (Fig. S5, see Supplementary Text for discussion).

Within the proteome, double cysteines are typically buried and highly conserved, participating in catalytic centers (e.g., thiol oxidoreductases) and structural domains (e.g., zinc fingers) (*6, 16*). In later sections, we explore how Cullin-RING ligases target misfolded zinc fingers proteins, suggesting that metal-mediated recognition of exposed cysteines – first characterized for FEM1B (*17, 19*) – represents a more general proteome surveillance strategy.

### Single cysteine degrons are degraded in a ubiquitin-independent manner

In contrast to the double cysteine degrons, the single cysteine degrons contained only one critical cysteine; additional cysteines in the peptide were not specifically required for degradation. The critical cysteine was usually flanked by a serine or threonine on its N-terminal side and an arginine on its C-terminal side (Fig. 3A-C, fig. S2I-K), although we identified several other variants of this motif, including NCR, HCR, TCA, and RCS (fig. S2L-P). In total, about 2-5 adjacent residues contributed to degradation and constituted the degron. Many of these residues are hydrophilic, suggesting that the recognition mechanism is not primarily driven by hydrophobicity. Glycine and proline were usually prohibited within this region, suggesting that the recognition for degradation may require a specific secondary structure such as an alpha helix.

Outside of this 3-6 amino acid degron, we noticed that bulky residues were sometimes required in the last five residues of the 34-residue peptides. This was most apparent for peptides LAMA1-26 and COL4A2-3 (fig. S2N-P), but was also seen for ELAPOR2-8 and C8A-3 (Fig. 3B, C). We speculate that these bulky C-terminal residues may help engagement by the proteasomal ATPase motor or mechanical unfolding of the peptide-fused GFP moiety, as discussed further below.

We performed CRISPR knockout screens on five single cysteine substrates, as in before, to identify the cognate E3 ligases. While we readily identified 26S proteasome subunits, no E2 conjugating enzymes or E3 ligases were recovered (fig. S4).

To circumvent the concern that relevant ubiquitylation enzymes might be unannotated and thus absent from our focused library, we next performed genome-wide CRISPR screens. These screens again robustly identified components of the 26S proteasome, but few other genes reached statistical significance. Some RNA processing genes such as XRN1 and the NELF complex were identified, but these have consistently scored in unrelated prior screens and are thus likely non-specific hits (*13, 21*).

Since our genome-wide screens did not reveal any putative E3 ubiquitin ligases or other ubiquitylation enzymes, the single cysteine substrates may be targeted to the proteasome in a ubiquitin-independent manner.

In light of these findings, we revisited our initial stability profiling screen with the goal of systematically classifying substrates based on their dependence on the ubiquitin-proteasome system for degradation. Using the library of cysteine-containing peptides described in Fig. 1A, we repeated the stability profiling assay in the presence of bortezomib (proteasome inhibition), TAK-243 (E1 activating enzyme inhibition), or MLN-4924 (Cullin-RING ligase inhibition) (Fig. 3G). Substrates fell into two major categories: those stabilized by bortezomib but unaffected by TAK-243 and MLN4924, suggesting ubiquitin-independent proteasomal degradation; and those stabilized by bortezomib, TAK-243, and MLN-4924, suggesting ubiquitin-dependent proteasomal degradation driven by Cullin-RING E3 ligase activity. Indeed, we were able to classify thousands of substrates as subject to either ubiquitin-independent or Cullin-dependent degradation (fig. S6A-E), including most of the top-ranked cysteine-dependent substrates (Fig. 3H). Strikingly, all of the substrates containing single cysteine degrons such as FAM155A-1 and ELAPOR2-8 underwent ubiquitin-independent degradation, while all the substrates with double cysteine degrons, including FBXO10-1, exhibited Cullin-RING ligase-dependent degradation. We were readily able to validate these conclusions in individual experiments (Fig. 3I). Altogether, these data define two broad classes of cysteine-dependent degradation: double-cysteine degrons targeted by a variety of Cullin-RING E3 ligases, and single-cysteine degrons degraded by the proteasome in a ubiquitin-independent manner.

### Mechanistic insights into ubiquitin-independent proteasomal degradation

Although the majority of proteasomal substrates require ubiquitylation for targeting, a handful of proteasomal substrates are degraded in a ubiquitin-independent manner. The most prominent example is ornithine decarboxylase (ODC), which in the late 1980’s became the first ubiquitin-independent substrate identified (*9*–*11, 22*). ODC, which catalyzes the rate-limiting step in polyamine synthesis, is regulated by a negative feedback loop in which high polyamine levels trigger exposure of a 37-amino-acid stretch at its C-terminus, resulting in proteasomal degradation (*23*). Although the underlying molecular mechanism by which ODC is recognized by the proteasome has not been resolved, the ODC C-terminus contains two elements critical for its degradation: the final five C-terminal residues, and, intriguingly, a central cysteine (*12*). Therefore, we hypothesized that the ubiquitin-independent degradation of ODC and our substrates bearing single-cysteine degrons may occur via a common proteasomal pathway.

We compared the degradation characteristics of ODC to that of our single cysteine substrates. Like our substrates, the ODC peptide is sufficient to induce cysteine-dependent, ubiquitin-independent proteasomal degradation, as assessed by flow cytometry (Fig. 3J). As a complementary way to assess protein stability, we used a cycloheximide chase assay. After blocking the production of new proteins with cycloheximide, we monitored protein decay over time through western blotting (Fig. 3K). We found that FAM155A-1 was rapidly degraded (t_1/2_ ≈ 5 mins) in a process dependent on its cysteine (abolished by cysteine-to-alanine substitution) and the proteasome (blocked by bortezomib), but not dependent on ubiquitylation (unaffected by TAK-243). The ODC C-terminal peptide behaved similarly and was rapidly degraded (t_1/2_ ≈ 5 mins) in a cysteine-dependent, proteasome-dependent, but ubiquitin-independent manner. In contrast, degradation of the double cysteine degron FBXO10-1 (t_1/2_ ≈ 15 mins) was entirely dependent on ubiquitylation.

We next asked whether an unstructured region is required for the degradation of single-cysteine substrates. All proteasomal substrates, whether targeted in a ubiquitin- or independent manner, are generally assumed to require a flexible initiation region that is long enough to pass through the static N-ring and reach into the ATPase ring of the proteasomal motor for engagement and mechanical translocation (*24*–*27*). These initiation regions often contain some bulky residues, which are readily engaged by the ATPase motor, whereas low-complexity sequences such as glycine-serine (GS) repeats can inhibit proteasomal degradation by disrupting ATPase engagement and mechanical force application (*28, 29*).

As GFP is highly structured, we hypothesized that the region C-terminal to the critical cysteine is acting as the initiation region to allow for efficient proteasomal degradation. This same region is also used as the initiation region for ODC (*12, 30*). Indeed, while various terminal sequences were permissive for the degradation of FAM155A-1 (fig. S6F, G), appending GS repeats inhibited its degradation (fig. S6H). We also investigated the effect of peptide length. Ubiquitin-independent degradation could occur with a C-terminal tail as short as 11 residues long, but a 6-residue tail abolished degradation (fig. S6I, J). This suggests that the ubiquitin-independent substrates are positioned especially close to the proteasomal ATPase ring, since only ∼11 residues (spanning ∼38Å in a fully extended conformation) are required. While our model substrates use the C-terminus as an initiation region, previous studies have shown that the N-terminus or internal loops can also serve as proteasomal initiation regions (*31*–*33*), and may be used by other substrates of this pathway.

### A CRISPR base editor screen identifies the proteasomal cysteine binding site

To elucidate the mechanism through which the proteasome recognizes exposed cysteine degrons, we sought to identify mutations in proteasome subunits that abrogated the degradation of substrates containing single-cysteine degrons. To achieve this, we conducted a base editor CRISPR screen (Fig. 4A); unlike standard CRISPR screens whose goal is complete disruption of gene function, base editor screens seek to introduce point mutations to enable fine-mapping of critical residues within essential proteins. Exploiting *Streptococcus pyogenes* Cas9 variants with relaxed PAM specificities, we designed a library of ∼20,000 sgRNAs tiling across 26 target genes, which included all components of the 19S regulatory particle plus the proteasome-associated factors MIDN, TXNL1, and PITHD1 (fig. S7A). This library was introduced into cells expressing the single cysteine ubiquitin-independent substrate ELAPOR2-8 in the GPS reporter system. We also introduced an adenine base editor that converts A•T to G•C within a ∼8-bp editing window (*34*). Mutant cells in which the reporter exhibited stabilization were isolated by FACS and sgRNA abundance was quantified by deep sequencing (fig. S7B).

**Fig 4.**
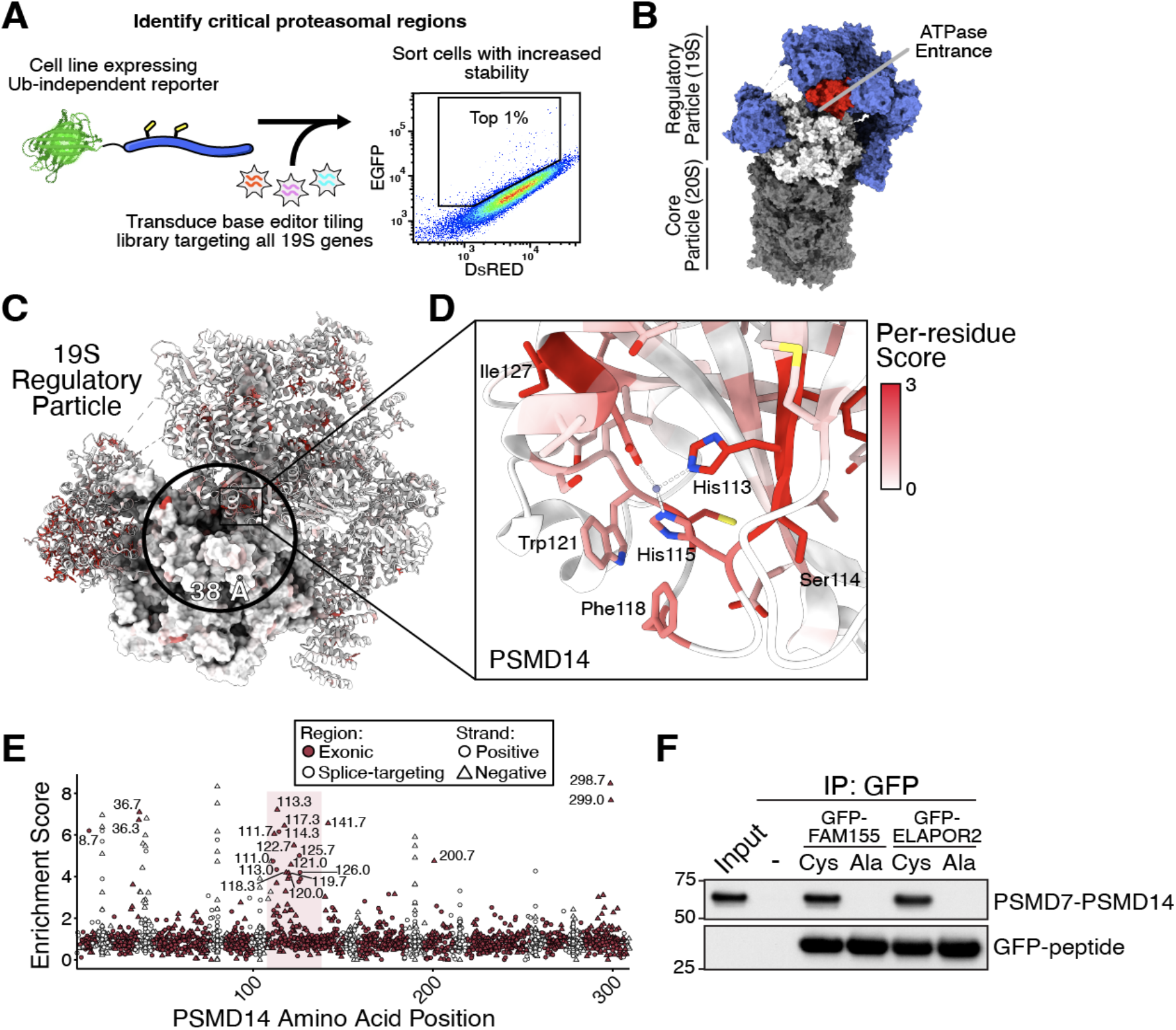
A base editor tiling screen identifies proteasomal determinants of ubiquitin-independent degradation. (**A**) A cell line expressing the ELAPOR2-8 stability reporter was transduced with a CRISPR base editor library targeting all subunits in the 19S regulatory particle as well as several proteasome-associated factors. Each guide included a linked reporter sequence, enabling direct identification of the mutations introduced by each guide. Cells with stabilized reporters were enriched and guides were sequenced. (**B**) The 19S regulatory particle is composed of the ATPase subunits (white), PSMD14 (red), and other non-ATPase subunits (blue) (PDB: 8USB) (*15*). (**C**) Residue-level functional scores from the screen were mapped onto the structure of the human 19S regulatory particle. ATPase hexamer is shown in surface representation, whereas other 19S subunits are shown in cartoon representation. A patch in PSMD14 was the highest scoring region within 38Å of the ATPase motor entrance. (**D**) Multiple residues at or near the active site of PSMD14 scored highly. (**E**) Enrichment scores for individual sgRNAs targeting PSMD14. Splice-disrupting guides are shown in gray, and the region encompassing the active site is shaded in light red. (**F**) A PSMD7–PSMD14 fusion construct was expressed in human cells and used in a pulldown assay to test its interaction with immobilized cysteine substrates. Input represents 0.5% of the total lysate; pulldown represents 100% of the bound fraction.

Because base editing outcomes can be variable and hard to predict computationally, we included a target reporter construct alongside each sgRNA (*35*). This reporter mirrored the endogenous sequence and underwent editing in parallel, enabling direct measurement of the specific mutations introduced by each guide. With this data, we created a custom pipeline that allowed us to resolve per-residue functional scores across the 19S regulatory particle with high resolution (fig. S7C). The screen exhibited good reproducibility across two replicates (fig. S8). Positive control guides (such as those targeting splice sites) typically abolished degradation, whereas negative control guides (such as those targeting intergenic regions) typically had little effect. Guides targeting genes encoding subunits of the 19S particle scored consistently, whereas those targeting accessory factors – including USP14, UCHL5, MIDN, TXNL1, and PITHD1 – did not (fig. S7B, S8), consistent with our prior CRISPR knockout screen data (Fig. 3D-F).

Numerous high-scoring residues were located within 19S subunits (Fig. 4B-D), indicating mutations that impaired degradation of our model single-cysteine degron substrate. Mapping those scores onto the structure of the 19S regulatory particle revealed many residues that were buried, which likely disrupt proteasome folding when mutated (fig. S7D), as well as residues at subunit interfaces (fig. S7E). However, since our prior experiments suggested that the engagement of cysteines may occur within ∼38Å of the proteasomal ATPase motor, we surveyed this region in detail and identified a site on PSMD14 as the leading candidate (Fig. 4C, D). PSMD14 (yeast Rpn11) is a JAMM-family metalloprotease that is responsible for deubiquitylating substrates as they are entering the proteasomal ATPase. Multiple guides targeting the catalytic region of PSMD14 scored, with consistent enrichment across replicates (Fig. 4E, fig. S7F).

To test whether PSMD14 can bind these substrates, we performed co-immunoprecipitation assays. Because PSMD14 is an obligate heterodimer (*36, 37*), we engineered a stable fusion of PSMD14 with its partner PSMD7, omitting the coiled-coil regions required for proteasome incorporation. When expressed in cells, this PSMD14–PSMD7 fusion selectively bound GFP fusion substrates exposing cysteine degrons, as confirmed by its co-immunoprecipitation (Fig. 4F). Together, these data illustrate the power of base editor screening for dissecting functional sites within essential multi-protein complexes and suggest that PSMD14 may function as a ubiquitin-independent receptor for exposed cysteine degrons.

### Single cysteines degrons are directly recognized by the proteasome deubiquitinase PSMD14/RPN11

PSMD14 is known to possess a low-affinity ubiquitin-binding site near its active site (*38*). Given the strong signal for PSMD14 in our base editor screen, we hypothesized that exposed degrons might directly compete with ubiquitin at this active site. To test this notion, we employed an *in vitro* assay to measure PSMD14 deubiquitinase activity towards fluorescently-labeled ubiquitin (*38, 39*). The cleavage of a fluorescently labeled ubiquitin by PSMD14 was measured through changes in fluorescence polarization, either in the absence or presence of cysteine-containing peptides (Fig. 5A). We purified recombinant PSMD14–PSMD7 heterodimer from *E. coli* and synthesized peptides harboring single cysteine degrons derived from three substrates (ODC, FAM155A, and ELAPOR2) that undergo ubiquitin-independent degradation, thereby ensuring a system free of eukaryotic cofactors. We found that each of the cysteine-containing peptides inhibited PSMD14 deubiquitinase activity with low micromolar IC_50_ values, consistent with direct competition at the active site, whereas control peptides with the critical cysteine mutated to alanine did not (Fig. 5, B and C). These findings could not be explained by nonspecific interactions with metalloenzymes, as the peptides had no effect on the activity of AMSH, a related JAMM-family deubiquitinase (fig. S9B) (*40*). Furthermore, the cysteine-containing peptides also inhibited the deubiquitinase activity of the yeast Rpn11–Rpn8 heterodimer, the yeast ortholog of human human PSMD14-PSMD7, suggesting conservation of the recognition mechanism from yeast to humans (Fig. 5, D and E).

**Fig 5.**
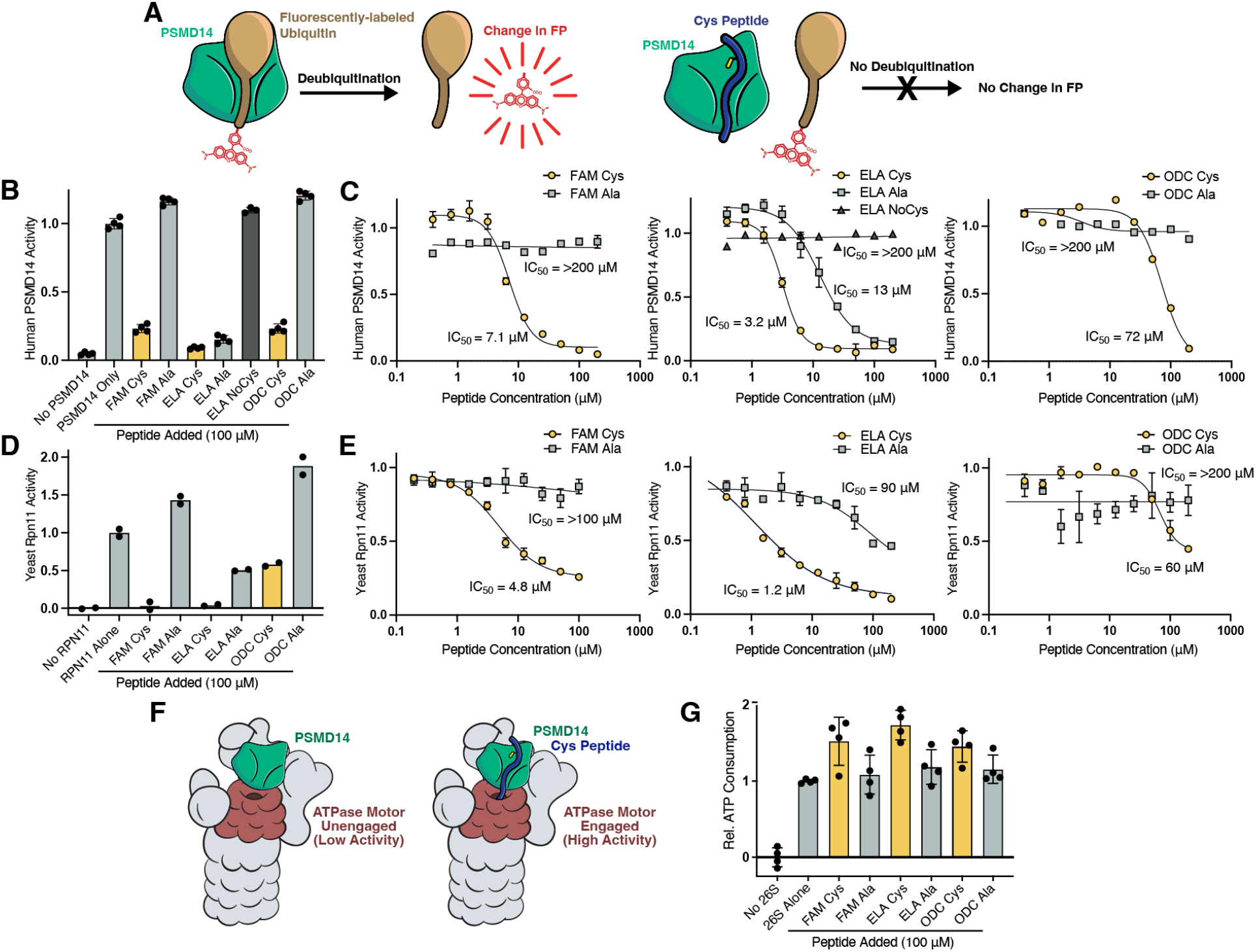
Cysteine substrates are engaged by the active site of PSMD14/Rpn11. (**A**) A fluorescence polarization (FP) assay with ubiquitin-TAMRA was used to measure the deubiquitinase activity of recombinant human PSMD14 and its yeast ortholog Rpn11. Liberation of TAMRA upon ubiquitin cleavage leads to a decrease in FP. (**B**) Cysteine-containing peptides inhibit PSMD14 activity, consistent with direct engagement of its catalytic site. Activity was measured in the presence of 100 μM synthetic peptides. (**C**) Dose-dependent inhibition of PSMD14 activity by cysteine-containing peptides. (**D, E**) Summary of yeast Rpn11 inhibition across multiple cysteine peptides, shown at a fixed concentration (100 μM) (D) and as a titration series (E). (**F**) Following substrate binding, engagement of the proteasomal ATPase motors enables unfolding and translocation into the proteolytic core, a step coupled to ATP hydrolysis. (**G**) Synthetic cysteine-containing peptides stimulate ATP consumption by reconstituted, partially recombinant yeast 26S proteasomes, indicating productive engagement of the ATPase motor. Alanine-substituted control peptides do not show this effect.

We next asked whether the interaction between the cysteine peptides and PSMD14 could involve disulfide bonding. A cysteine-less version of yeast Rpn11–Rpn8, in which all cysteine residues were substituted with alanine, retained wild-type enzymatic activity and remained sensitive to inhibition by the cysteine-containing peptides (fig. S9C), demonstrating that substrate binding does not depend on intermolecular disulfide bonds. After initial binding, substrates must also engage the proteasomal ATPase motor to be unfolded and translocated into the proteolytic core for degradation. Cysteine-containing peptides, but not their alanine-substituted controls, caused a 50–70% increase in ATP hydrolysis by yeast 26S proteasomes (Fig. 5G), suggesting that these substrates productively engage the ATPase motor to support their active processing by the proteasome. Altogether, these results demonstrate sequence-specific recognition of cysteine degrons by PSMD14/Rpn11, leading to substrate engagement and activation of the ATPase motor independent of ubiquitin.

### Protein quality control via cysteine degrons involves both ubiquitin-dependent and -independent pathways

These observations led us to hypothesize that the ubiquitin-independent cysteine recognition might play a role in degrading misfolding cysteine-containing proteins, such as those with zinc finger domains. Zinc fingers are among the most abundant cysteine-rich domains in eukaryotes and are central to transcriptional regulation, ubiquitylation, and apoptosis (34). In their native state, zinc finger cysteines are buried around a coordinated zinc ion; upon unfolding, however, exposure of these cysteines might render the polypeptide susceptible to proteasomal degradation.

Intriguingly, our saturation mutagenesis experiments (Fig. 3A-C, fig. S2J, K, O) suggested that the introduction of additional cysteine or histidine residues downstream of cysteine degrons occasionally led to increased stability. This effect was most pronounced for the GFP fusion with the C8A-3 peptide (Fig. 6A), but a weak effect could also be observed with ELAPOR2-8 and FAM155A-1 (Fig. 3A, B). In combination with the native cysteines, we hypothesized that an extra cysteine or histidine residue might allow for the coordination of a zinc ion; the resulting zinc finger-like structure would therefore shield the critical cysteines from recognition by the proteasome (Fig. 6B). We began to test this model by examining the C8A-3 peptide substrate. Two mutations, G14C and Q19H, predicted by AlphaFold 3 to result in zinc-binding pockets, conferred a substantial stabilizing effect (Fig. 6C-D). Treatment with the cell-permeable zinc chelator TPEN reversed this effect, but had no impact on the stability of the wild-type substrate or a C4A mutant lacking the critical cysteine (Fig. 6D). Thus, zinc coordination can shield cysteine degrons from ubiquitin-independent degradation by the proteasome.

**Fig 6.**
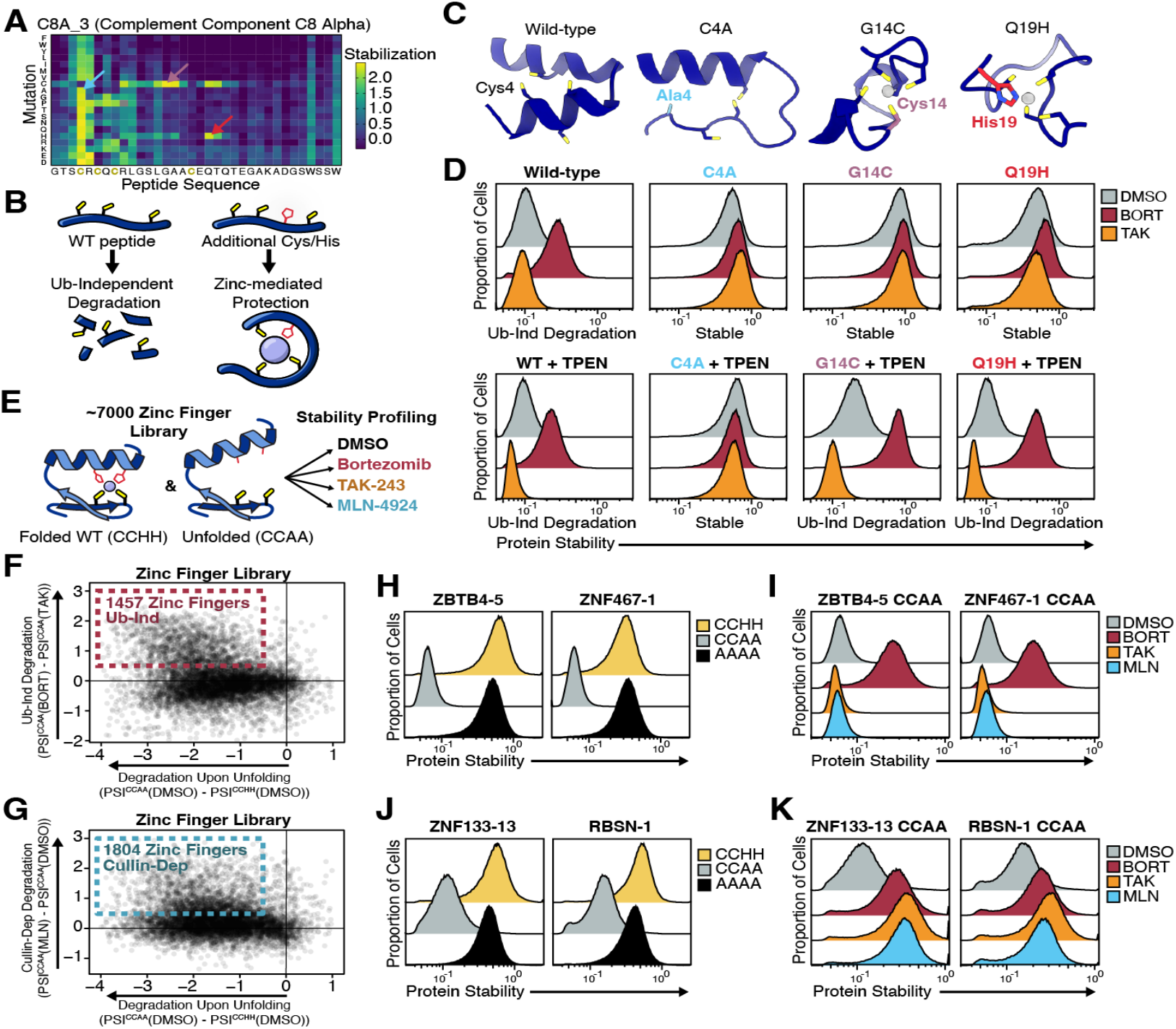
Both cysteine surveillance pathways target unfolded zinc finger domains. (**A**) Saturation mutagenesis heatmap for the C8A-3 peptide, showing that introduction of cysteine or histidine residues at specific positions can block degradation. (**B**) The wild-type peptide contains an exposed cysteine that is recognized by the proteasome. Mutations introducing additional cysteines or histidines may coordinate zinc, forming a folded structure that occludes the reactive cysteine. (**C**) AlphaFold 3 structural predictions of wild-type C8A-3 and three degradation-resistant mutants. The G14C and Q19H variants are predicted to form zinc-binding motifs resembling zinc fingers. (**D**) Stability of C8A-3 and its mutants was assessed in cells treated with bortezomib or TAK-243. Addition of the zinc chelator TPEN restored ubiquitin-independent degradation of the G14C and Q19H mutants. (**E**) A GPS-based stability screen of ∼7,000 C2H2-type zinc finger peptides, comparing wild-type (CCHH) versus mutant (CCAA) versions that simulate domain unfolding. Inhibitor treatments were used to classify degradation pathways. (**F, G**) Each zinc finger was evaluated for ubiquitin-independent degradation upon unfolding (F) or Cullin-dependent degradation upon unfolding (G). (**H, I**) Representative zinc fingers identified in (F) were tested for cysteine-dependent degradation after unfolding (H) and for proteasome-dependent, ubiquitin-independent degradation (I). (**J, K**) Representative zinc fingers from (G) were similarly evaluated for cysteine-dependence (J) and for Cullin-dependent degradation (K) upon unfolding.

To explore the generalizability of these observations, we performed a high-throughput stability screen to systematically examine ∼7,000 C2H2 zinc finger domains (Fig. 6E). For each domain we engineered a mutant in which both histidine residues were replaced with alanine (CCHH → CCAA), thus disrupting zinc coordination and promoting unfolding. Strikingly, zinc finger unfolding caused increased degradation for 5,985 domains (86%), with 3,497 domains (50%) exhibiting a very substantial effect (>1.5 PSI units) (fig. S10A). Stability profiling in the presence of inhibitors demonstrated that 1,457 domains (21%) were degraded in a ubiquitin-independent, proteasome-dependent manner (Fig. 6F), while 1,804 (26%) underwent ubiquitin-mediated degradation driven by Cullin-RING E3 ligases (Fig. 6G). We validated these results using representative zinc fingers from each class. The ubiquitin-independent cases required exposed cysteines for degradation to occur and were insensitive to inhibition of ubiquitylation (Fig. 6H–I, fig. S10B–E); conversely, Cullin-dependent domains also exhibited cysteine-dependent degradation but required components of the ubiquitylation machinery (Fig. 6J–K, fig. S10F–G). Together, these findings demonstrate that both cysteine-directed degradation pathways – ubiquitin-independent and Cullin-RING ligase-dependent – contribute to the surveillance of misfolded zinc fingers.

### Cysteine-directed ubiquitin-independent degradation regulates multiple endogenous proteins

Next, we sought to identify endogenous proteins regulated in a ubiquitin-independent manner via cysteine degrons. To do this we performed whole-cell proteomics in three cell lines, seeking proteins whose abundance was increased by the proteasome inhibitor bortezomib but not by the ubiquitin E1 inhibitor TAK-243. High-confidence candidates meeting these criteria, included known substrates of ubiquitin-independent degradation including ODC and FOS, as well as the proteasome cofactors ZFAND5, AKIRIN2, and MIDN (Fig. 7A-C). We also uncovered new candidate substrates, most notably OSER1, which was among the top hits in two cell lines. These proteins were robustly stabilized by proteasome inhibition but unaffected by E1 or Cullin-RING ligase inhibition (fig. S11A).

**Fig 7.**
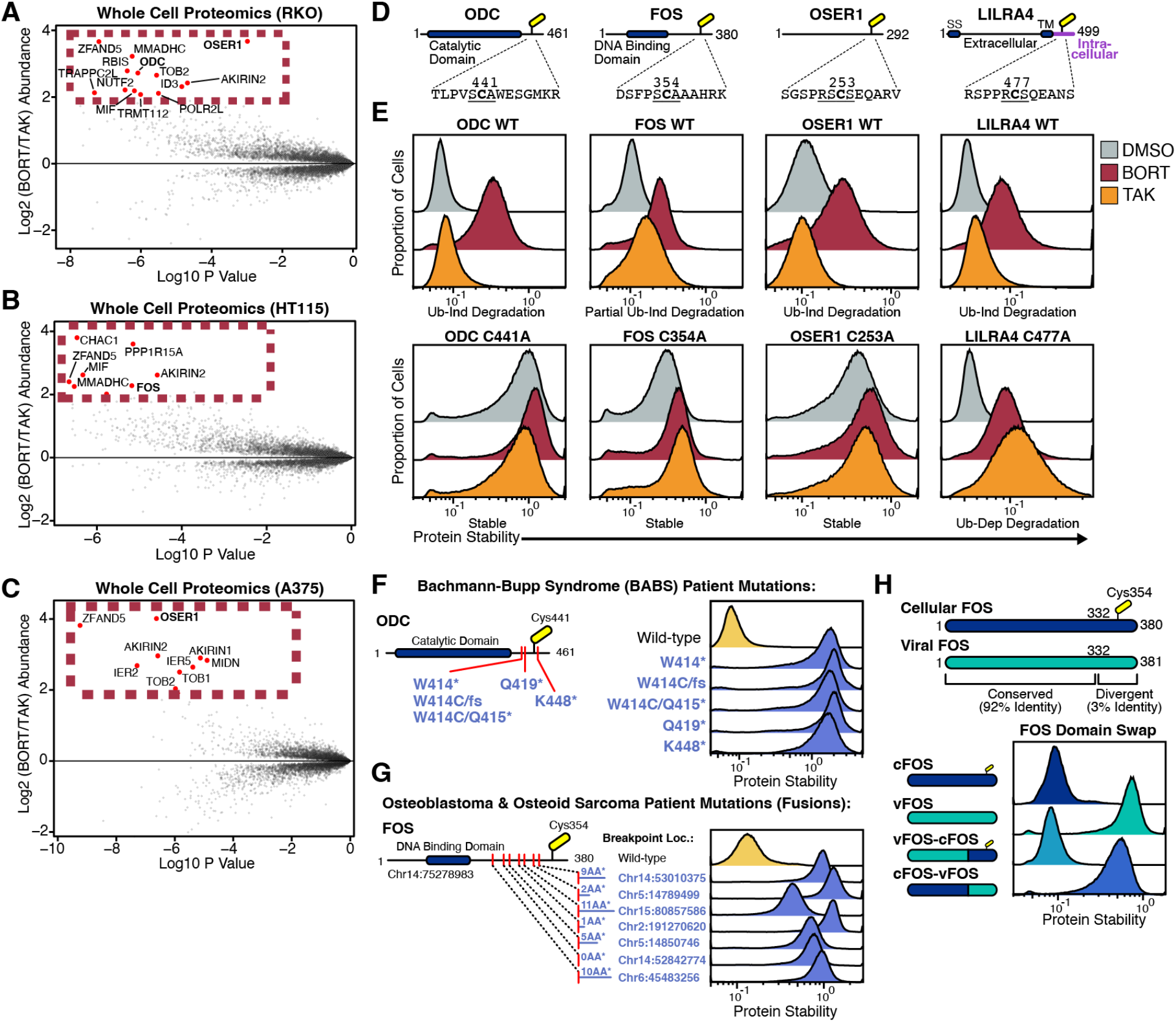
Endogenous substrates of the ubiquitin-independent cysteine pathway. (**A**-**C**) Whole-cell proteomics was performed on three cell lines: RKO (A), HT115 (B), and A375 (C). Volcano plots comparing protein abundances in the presence of bortezomib versus TAK-243 are shown. (**D**) Schematics of the domain architecture of ubiquitin-independent substrates ODC, FOS, OSER1, and LILRA4 are shown, with key cysteines indicated. (**E**) Protein stability reporter assays were performed on wild-type substrate, as well as cysteine-to-alanine mutants, and assessed in the presence of bortezomib or TAK-243. (**F**) Point mutations in ODC associated with Bachmann-Bupp Syndrome (left) were introduced and evaluated for effects on protein stability (right). (**G**) FOS mutations identified in patients with osteoblastoma and osteoid sarcoma were similarly tested for altered stability. (**H**) Domain-swap experiments between cellular FOS and viral FOS (from FBJ murine osteosarcoma virus) demonstrate that the C-terminal region regulates protein stability.

To validate these findings, we examined the stability of full-length ODC, FOS, and OSER1 in the context of the GPS system (Fig. 7D-E). All three proteins were degraded in a ubiquitin-independent manner. Like its C-terminal peptide (Fig. 3J), degradation of full-length ODC was entirely dependent on a single cysteine residue (Cys-441) located within an SCA motif. FOS degradation was partially independent of ubiquitylation, and also required a conserved cysteine (Cys-354) in an SCA motif (Fig. 7E). Mutation of this cysteine to alanine (C354A) increased FOS stability 2.7-fold and blocked ubiquitin-independent degradation. Indeed, fusing the C-terminal residues of FOS peptide to GFP was sufficient to confer cysteine-dependent, ubiquitin-independent degradation (fig. S11B), and we identified the four C-terminal residues (LLAL) of FOS as a likely initiation motif (fig. S11C). Lastly, by systematically mutating each of its cysteine residues, we determined that ubiquitin-independent degradation of OSER1 requires Cys-253. Given the role of OSER1 in oxidative stress responses (*41, 42*), it will be important to determine whether its degradation is regulated by cellular stress.

We also re-examined LILRA4, a transmembrane protein previously identified via GPS reporters as a ubiquitin-independent substrate (*14*). The ubiquitin-independent degradation of LILRA4 is also entirely dependent on a single cysteine (Cys-477) (Fig. 7E). Intriguingly, LILRA4 is only degraded in a ubiquitin-independent manner when its N-terminal signal sequence is disrupted by fusion to GFP; when the N-terminal signal sequence is intact, allowing for proper localization to the membrane (fig. S11E), LILRA4 is not degraded (fig. S11E, F). This further supports that the ubiquitin-independent cysteine pathway is attuned to recognizing aberrant proteins.

### Disease-linked mutations impair cysteine-directed proteasomal degradation

Finally, we considered the physiological consequences resulting from dysregulation of ubiquitin-independent degradation via cysteine degrons. ODC is the rate-limiting enzyme in polyamine biosynthesis and, as excess polyamines are cytotoxic (*43*), is tightly regulated. Mutations that truncate the C-terminus of ODC cause the neurodevelopmental disorder Bachmann-Bupp Syndrome (BABS) (*44, 45*). Strikingly, all five of the BABS patient mutations abolished ODC degradation, suggesting that degradation of the cysteine degron underlies the molecular pathology of BABS.

FOS, a proto-oncogenic transcription factor, is recurrently activated in osteoblastoma and osteoid sarcoma through gene fusions that truncate its C-terminus and replace it with a short (≤11 amino acids) segments from various partners (Fig. 7G) (*46, 47*). All seven tested fusion proteins derived from osteoblastoma and osteoid sarcoma patients exhibited marked stabilization; furthermore, many of these fusions stabilized FOS more strongly than the C354A mutation alone (Fig. 7E), suggesting that they may disrupt both the cysteine-directed and canonical ubiquitin-mediated degradation.

Cysteine degrons have also been subverted by retroviruses. The Finkel-Biskis-Jinkins (FBJ) murine osteosarcoma virus encodes vFOS, one of the earliest described oncogenes (*48, 49*). Although vFOS is highly similar to cellular FOS (cFOS), it harbors a frameshift at the C-terminus that replaces the region containing the cysteine degron with an unrelated 49-residue tail (*50*). Domain-swapping experiments between vFOS and cFOS revealed that the C-terminal tail is both necessary and sufficient for degradation (Fig. 7H, fig. S11G), indicating that loss of this degron may be a key oncogenic mechanism shared by viral and tumor-derived FOS variants. Importantly, all these pathogenic mutations and truncations in ODC and FOS occurred downstream of their enzymatic or DNA-binding domains, preserving their functional activity but impairing their degradation. Altogether, these data demonstrate that ubiquitin-independent degradation via cysteine degrons is critical for cellular homeostasis, with mutational escape from this pathway resulting in disease.

## DISCUSSION

Protein homeostasis is fundamental to cellular health, and its disruption underlies a wide range of human diseases. To maintain proteome integrity, cells rely on surveillance systems that detect and eliminate misfolded or aberrant proteins, typically by tagging them with ubiquitin and targeting them to the 26S proteasome. These established quality control pathways recognize features such as exposed hydrophobic residues or orphaned subunits from failed complex assembly (*51*–*53*). Here, we find that exposed cysteines also serve as a general degradation signal, engaging two distinct proteasomal pathways. One involves Cullin-RING E3 ligases that detect paired cysteines and promote ubiquitin-dependent degradation. The other is a ubiquitin-independent pathway in which the 26S proteasome directly engages exposed cysteines via its PSMD14/Rpn11 subunit. The convergence of distinct mechanisms highlights the strong evolutionary pressure to monitor and eliminate proteins harboring reactive cysteine residues.

Our genetic and biochemical data suggest that single cysteine substrates are directly recognized by the 26S proteasome. In particular, synthetic cysteine-containing peptides can inhibit purified PSMD14 and can stimulate 26S proteasome in reconstituted systems free from contaminating factors. The direct engagement of cysteines is distinguished from previously described ubiquitin-independent mechanisms that rely on ubiquitin-like domains and modifiers including Midnolin, Ubiquilin, and Fat10 (*13*–*15*). While cysteine substrates can be directly bound to PSMD14, we note that ODC antizyme (OAZ1) is required for efficient degradation of full-length ODC (*11, 54*). It remains possible that analogous factors could modulate the degradation of cysteine-containing substrates in vivo, in a manner not detectable through current genetic screening technology. Likewise, environmental triggers such as redox stress or zinc depletion might dynamically regulate these pathways, and warrants further investigation.

Our findings also resolve a longstanding puzzle: why the proteasome evolved a specialized ubiquitin-independent mechanism to degrade a seemingly singular substrate, ornithine decarboxylase (ODC) (*55, 56*). Although yeast lack the specific ODC degron (*57, 58*), their proteasomes can still degrade human ODC (*59*–*61*), suggesting the existence of a conserved ubiquitin-independent degradation pathway. We propose that direct recognition of exposed cysteines by PSMD14/Rpn11 evolved early in eukaryotic history as a general quality control mechanism, and that key vertebrate proteins, including ODC, FOS, and OSER1, later co-opted this system to enable their own rapid, regulated degradation.

Beyond basic biology, our findings have clear relevance to disease. Mutations that disrupt the cysteine degron of ODC stabilize it and cause Bachmann-Bupp Syndrome. Similarly, gene fusions in osteoblastoma frequently remove the degron-bearing C-terminus of FOS, thereby converting it into a hyperstable oncogene. Intriguingly, this mechanism is recapitulated by vFOS, a retroviral oncogene that achieves oncogenic stabilization by replacing its degron with a non-degradable tail lacking the critical cysteine.

Multiple pathways have converged to regulate the proto-oncogene FOS. In addition to the direct cysteine-dependent pathway described here, we previously showed that FOS is targeted by the Midnolin-mediated ubiquitin-independent pathway (*13*), and it can also be targeted by canonical ubiquitin-dependent degradation (*8, 62*). Our data indicate that in HEK293T cells, the cysteine degradation pathway is critical for the ubiquitin-independent degradation of FOS; however in alternative conditions, such as in neurons with increased Midnolin expression, the Midnolin degradation pathway may play a more prominent role. The presence of multiple, overlapping degradation pathways likely reflects the need for tight regulation of FOS activity, guarding against aberrant expression that could drive tumorigenesis.

Why does the proteasome selectively recognize certain cysteines but not others? Our data underscore the importance of adjacent unstructured regions (fig. S6), suggesting that this pathway preferentially targets unfolded or misfolded proteins. In addition, we observe that cysteine recognition is more likely at the N-terminal side of an α-helix and in proximity to arginine or threonine residues (Fig. 3A-C, fig. S2I-P). These structural features are known to stabilize the cysteine thiolate form by lowering its pKa, and increase its propensity for oxidative modification (*4, 63*–*65*). The proteasome may thus be tuned to recognize cysteines with heightened chemical reactivity.

Taken together, our work defines cysteines as a degradation signal and advances our understanding of how cells integrate structural integrity with degradative fate. Further mechanistic insights will be critical to fully elucidate these pathways, and they may enable new avenues for therapeutics such as molecular glues that bypass ubiquitin and guide substrates directly to the proteasome.

## Acknowledgements

We would like to thank Kwangwoon Lee and Sam Whedon for assistance with biochemistry; Maheeshi Abeywardana for assistance with mass spectrometry; Aoyue Mao and Anthony Liang for assistance with cellular assays; Matthew Francoeur, Quang Vinh Phan, Tian Yu, and Jayoung Ryu for assistance with base editor screening; Dan Finley, Bryan Seguinot, Alfred Goldberg, Donghoon Lee, and Cynthia Morton for reagents; Ellen Shrock and Caleb Glassman for critically reading the manuscript; and Phil Cole, Kim Orth, and all members of the Elledge laboratory for helpful suggestions.

## Author Contributions

E.M. performed genetic screens; K.C.D. performed in vitro experiments; E.M., A.K., and M.S. performed cellular assays; K.A.D., H.-T.H., and B.Y. performed proteomics under the supervision of E.S.F.; R.L.S. assisted with the design and analysis of CRISPR base editor screens; R.T.T. designed GPS libraries; A.M. and S.J.E. supervised the study; E.M., R.T.T., A.M., and S.J.E wrote the manuscript.

## Data and material availability

Any reagents that are unique to this study will be made available upon request.

