## Supplementary Data for "Proteasomal surveillance of exposed cysteines"

#### Materials and Methods

Cell culture

All screens and protein stability reporter experiments were performed in HEK-293T/17 cells (CRL-11268) obtained from ATCC. Approximately every 6 months, cells were checked to be free of mycoplasma using the MycoAlert Detection Kit (Lonza). HEK-293/17 cells were grown in DMEM supplemented with 10% fetal bovine serum (FBS) and 1% penicillin-streptomycin (Gibco), RKO cells (ATCC) were grown in EMEM supplemented with 10% FBS, HT115 cells were grown in DMEM supplemented with 15% FBS, and A375 cells (ATCC) were grown in DMEM supplemented with 10% FBS.

Constructs

Gene fragments encoding individual peptides – including zinc finger domains and C-terminal ODC and FOS fragments – were synthesized by Twist and inserted into GPS3.0-I24-Hygro using Hifi DNA Assembly (New England Biolabs).
Full-length ORFs were obtained from the Ultimate ORFeome collection (Thermo Fisher), or were synthesized by Twist Bioscience. They were introduced into GPS3.0-Hygro using LR Clonase II (Thermo Fisher) or Hifi DNA Assembly (NEB).

Single point mutations were introduced using Quikchange Site-Directed Mutagenesis (Agilent). More complex mutations, including multiple point mutations and fusions, were generated by synthesizing gene fragments (Twist Bioscience) followed by Hifi DNA Assembly (New England Biolabs). The 3xFLAG-mTagBFP2-PSMD7(1-182)-PSMD14(26-221) construct was cloned into pMax GFP (Lonza Bioscience) with the GFP removed. SS-2xHA-LILRA4 was cloned into pHAGE SFFV. The following reporter sequences were used (linker sequences between GFP and protein of interest are underlined):

*GPS-Peptide ("X" encodes peptide sequence):*
MVSKGEELFTGVVPILVELDGDVNGHKFSVSGEGEGDATYGKLTLKFICTTGKLPVPWP
TLVTTLTYGVQCFSRYPDHMKQHDFFKSAMPEGYVQERTIFFKDDGNYKTRAEVKFEG
DTLVNRIELKGIDFKEDGNILGHKLEYNYNSHNVYIMADKQKNGIKVNFKIRHNIEDGSV
QLADHYQQNTPIGDGPVLLPDNHYLSTQSALSKDPNEKRDHMVLLEFVTAAGITLGMD
ELYKGLATLRESGPIYFERIPLRXXXXXXXXXXXXXXXXXXXXXXXXXXXXXXXXXXX
*

*GPS-vFOS:*
MVSKGEELFTGVVPILVELDGDVNGHKFSVSGEGEGDATYGKLTLKFICTTGKLPVPWP
TLVTTLTYGVQCFSRYPDHMKQHDFFKSAMPEGYVQERTIFFKDDGNYKTRAEVKFEG
DTLVNRIELKGIDFKEDGNILGHKLEYNYNSHNVYIMADKQKNGIKVNFKIRHNIEDGSV
QLADHYQQNTPIGDGPVLLPDNHYLSTQSALSKDPNEKRDHMVLLEFVTAAGITLGMD
ELYKGLATLRESGPIYFETSLYKKAGTMMFSGFNADYEASSFRCSSASPAGDSLSYYHSP
ADSFSSMGSPVNTQDFCADLSVSSANFIPTVTATSTSPDLQWLVQPTLVSSVAPSQTRAP
HPYGLPTQSAGAYARAEMVKTVSGGRAQSIGRRGKVEQLSPEEEEKRRIRRERNKMAA
AKCRNRRRELTDTLQAETDQLEDKKSALQTEIANLLKEKEKLEFILAAHRPACKIPDDLG
FPEEMSVASLDLTGGLPEASTPESEEAFTLPLLNDPEPKPSLEPVKSISNVELKAEPFDDFL
FPASSRPSGSETSRSVPNVDLSGSFYAADWEPLHSNSLGMGPMVTELEPLCTPVVTCTPL
LRLPELTHAAGPVSSQRRQGSRHPDVPLPELVHYREEKHVFPQRFPST*

*GPS–FOS (cFOS):*
MVSKGEELFTGVVPILVELDGDVNGHKFSVSGEGEGDATYGKLTLKFICTTGKLPVPWP
TLVTTLTYGVQCFSRYPDHMKQHDFFKSAMPEGYVQERTIFFKDDGNYKTRAEVKFEG
DTLVNRIELKGIDFKEDGNILGHKLEYNYNSHNVYIMADKQKNGIKVNFKIRHNIEDGSV
QLADHYQQNTPIGDGPVLLPDNHYLSTQSALSKDPNEKRDHMVLLEFVTAAGITLGMD
ELYKGLATLRESGPIYFETSLYKKAGTMMFSGFNADYEASSSRCSSASPAGDSLSYYHSP
ADSFSSMGSPVNAQDFCTDLAVSSANFIPTVTAISTSPDLQWLVQPALVSSVAPSQTRAP
HPFGVPAPSAGAYSRAGVVKTMTGGRAQSIGRRGKVEQLSPEEEEKRRIRRERNKMAA
AKCRNRRRELTDTLQAETDQLEDEKSALQTEIANLLKEKEKLEFILAAHRPACKIPDDLG
FPEEMSVASLDLTGGLPEVATPESEEAFTLPLLNDPEPKPSVEPVKSISSMELKTEPFDDFL
FPASSRPSGSETARSVPDMDLSGSFYAADWEPLHSGSLGMGPMATELEPLCTPVVTCTPS
CTAYTSSFVFTYPEADSFPSCAAAHRKGSSSNEPSSDSLSSPTLLAL*

*FOS Chr14:53010375 (PD7521):*
MVSKGEELFTGVVPILVELDGDVNGHKFSVSGEGEGDATYGKLTLKFICTTGKLPVPWP
TLVTTLTYGVQCFSRYPDHMKQHDFFKSAMPEGYVQERTIFFKDDGNYKTRAEVKFEG
DTLVNRIELKGIDFKEDGNILGHKLEYNYNSHNVYIMADKQKNGIKVNFKIRHNIEDGSV
QLADHYQQNTPIGDGPVLLPDNHYLSTQSALSKDPNEKRDHMVLLEFVTAAGITLGMD
ELYKGLATLRESGPIYFETSLYKKAGTMMFSGFNADYEASSSRCSSASPAGDSLSYYHSP
ADSFSSMGSPVNAQDFCTDLAVSSANFIPTVTAISTSPDLQWLVQPALVSSVAPSQTRAP
HPFGVPAPSAGAYSRAGVVKTMTGGRAQSIGRRGKVEQLSPEEEEKRRIRRERNKMAA
AKCRNRRRELTDTLQAETDQLEDEKSALQTEIANLLKEKEKLEFILAAHRPACKIPDDLG
FPEEMSVASLDLTGGLPEVATPESEEAFTLPLLNDPEPKPSVEPVKSISSMELKTEPFDDFL
FPASSRPSGSETARSVDISSVFKVV*

*FOS Chr5:14789499 (FOS-ANKH):*
MVSKGEELFTGVVPILVELDGDVNGHKFSVSGEGEGDATYGKLTLKFICTTGKLPVPWP
TLVTTLTYGVQCFSRYPDHMKQHDFFKSAMPEGYVQERTIFFKDDGNYKTRAEVKFEG
DTLVNRIELKGIDFKEDGNILGHKLEYNYNSHNVYIMADKQKNGIKVNFKIRHNIEDGSV
QLADHYQQNTPIGDGPVLLPDNHYLSTQSALSKDPNEKRDHMVLLEFVTAAGITLGMD
ELYKGLATLRESGPIYFETSLYKKAGTMMFSGFNADYEASSSRCSSASPAGDSLSYYHSP
ADSFSSMGSPVNAQDFCTDLAVSSANFIPTVTAISTSPDLQWLVQPALVSSVAPSQTRAP
HPFGVPAPSAGAYSRAGVVKTMTGGRAQSIGRRGKVEQLSPEEEEKRRIRRERNKMAA
AKCRNRRRELTDTLQAETDQLEDEKSALQTEIANLLKEKEKLEFILAAHRPACKIPDDLG
FPEEMSVASLDLTGGLPEVATPESEEAFTLPLLNDPEPKPSVEPVKSISSMELKTEPFDDFL
FPASSRPSGSETGF*

*FOS Chr15:80857586 (PD13481):*
MVSKGEELFTGVVPILVELDGDVNGHKFSVSGEGEGDATYGKLTLKFICTTGKLPVPWP
TLVTTLTYGVQCFSRYPDHMKQHDFFKSAMPEGYVQERTIFFKDDGNYKTRAEVKFEG
DTLVNRIELKGIDFKEDGNILGHKLEYNYNSHNVYIMADKQKNGIKVNFKIRHNIEDGSV
QLADHYQQNTPIGDGPVLLPDNHYLSTQSALSKDPNEKRDHMVLLEFVTAAGITLGMD
ELYKGLATLRESGPIYFETSLYKKAGTMMFSGFNADYEASSSRCSSASPAGDSLSYYHSP
ADSFSSMGSPVNAQDFCTDLAVSSANFIPTVTAISTSPDLQWLVQPALVSSVAPSQTRAP
HPFGVPAPSAGAYSRAGVVKTMTGGRAQSIGRRGKVEQLSPEEEEKRRIRRERNKMAA
AKCRNRRRELTDTLQAETDQLEDEKSALQTEIANLLKEKEKLEFILAAHRPACKIPDDLG

FPEEMSVASLDLTGGLPEVATPESEEAFTLPLLNDPEPKPSVEPVKSISSMELKTEPFDDFL
KFANFTVHISH*

*FOS Chr2:191270620 (PD13482):*
MVSKGEELFTGVVPILVELDGDVNGHKFSVSGEGEGDATYGKLTLKFICTTGKLPVPWP
TLVTTLTYGVQCFSRYPDHMKQHDFFKSAMPEGYVQERTIFFKDDGNYKTRAEVKFEG
DTLVNRIELKGIDFKEDGNILGHKLEYNYNSHNVYIMADKQKNGIKVNFKIRHNIEDGSV
QLADHYQQNTPIGDGPVLLPDNHYLSTQSALSKDPNEKRDHMVLLEFVTAAGITLGMD
ELYKGLATLRESGPIYFETSLYKKAGTMMFSGFNADYEASSSRCSSASPAGDSLSYYHSP
ADSFSSMGSPVNAQDFCTDLAVSSANFIPTVTAISTSPDLQWLVQPALVSSVAPSQTRAP
HPFGVPAPSAGAYSRAGVVKTMTGGRAQSIGRRGKVEQLSPEEEEKRRIRRERNKMAA
AKCRNRRRELTDTLQAETDQLEDEKSALQTEIANLLKEKEKLEFILAAHRPACKIPDDLG
FPEEMSVASLDLTGGLPEVATPESEEAFTLPLLNDPEPKPSVEPVKSISSMELKTEPFDE*

*FOS Chr5:14850746 (PD13480):*
MVSKGEELFTGVVPILVELDGDVNGHKFSVSGEGEGDATYGKLTLKFICTTGKLPVPWP
TLVTTLTYGVQCFSRYPDHMKQHDFFKSAMPEGYVQERTIFFKDDGNYKTRAEVKFEG
DTLVNRIELKGIDFKEDGNILGHKLEYNYNSHNVYIMADKQKNGIKVNFKIRHNIEDGSV
QLADHYQQNTPIGDGPVLLPDNHYLSTQSALSKDPNEKRDHMVLLEFVTAAGITLGMD
ELYKGLATLRESGPIYFETSLYKKAGTMMFSGFNADYEASSSRCSSASPAGDSLSYYHSP
ADSFSSMGSPVNAQDFCTDLAVSSANFIPTVTAISTSPDLQWLVQPALVSSVAPSQTRAP
HPFGVPAPSAGAYSRAGVVKTMTGGRAQSIGRRGKVEQLSPEEEEKRRIRRERNKMAA
AKCRNRRRELTDTLQAETDQLEDEKSALQTEIANLLKEKEKLEFILAAHRPACKIPDDLG
FPEEMSVASLDLTGGLPEVATPESEEAFTLPLLNDPEPKPSVEPVKSICPLLN*

*FOS Chr14:52842774 (PD7519):*
MVSKGEELFTGVVPILVELDGDVNGHKFSVSGEGEGDATYGKLTLKFICTTGKLPVPWP
TLVTTLTYGVQCFSRYPDHMKQHDFFKSAMPEGYVQERTIFFKDDGNYKTRAEVKFEG
DTLVNRIELKGIDFKEDGNILGHKLEYNYNSHNVYIMADKQKNGIKVNFKIRHNIEDGSV
QLADHYQQNTPIGDGPVLLPDNHYLSTQSALSKDPNEKRDHMVLLEFVTAAGITLGMD
ELYKGLATLRESGPIYFETSLYKKAGTMMFSGFNADYEASSSRCSSASPAGDSLSYYHSP
ADSFSSMGSPVNAQDFCTDLAVSSANFIPTVTAISTSPDLQWLVQPALVSSVAPSQTRAP
HPFGVPAPSAGAYSRAGVVKTMTGGRAQSIGRRGKVEQLSPEEEEKRRIRRERNKMAA
AKCRNRRRELTDTLQAETDQLEDEKSALQTEIANLLKEKEKLEFILAAHRPACKIPDDLG
FPEEMSVASLDLTGGLPEVATPESEEAFTLPLLNDPEPKPSVEPV*

*FOS Chr6:45483256 (FOS-RUNX2):*
MVSKGEELFTGVVPILVELDGDVNGHKFSVSGEGEGDATYGKLTLKFICTTGKLPVPWP
TLVTTLTYGVQCFSRYPDHMKQHDFFKSAMPEGYVQERTIFFKDDGNYKTRAEVKFEG
DTLVNRIELKGIDFKEDGNILGHKLEYNYNSHNVYIMADKQKNGIKVNFKIRHNIEDGSV
QLADHYQQNTPIGDGPVLLPDNHYLSTQSALSKDPNEKRDHMVLLEFVTAAGITLGMD
ELYKGLATLRESGPIYFETSLYKKAGTMMFSGFNADYEASSSRCSSASPAGDSLSYYHSP
ADSFSSMGSPVNAQDFCTDLAVSSANFIPTVTAISTSPDLQWLVQPALVSSVAPSQTRAP
HPFGVPAPSAGAYSRAGVVKTMTGGRAQSIGRRGKVEQLSPEEEEKRRIRRERNKMAA
AKCRNRRRELTDTLQAETDQLEDEKSALQTEIANLLKEKEKLEFILAAHRPACKIPDDLG
FPEEMSVASLDLTGGLPEVATPESEEAFTLPLLNDPVQRSHTTNDC*

*GPS-LILRA4:*
MVSKGEELFTGVVPILVELDGDVNGHKFSVSGEGEGDATYGKLTLKFICTTGKLPVPWP
TLVTTLTYGVQCFSRYPDHMKQHDFFKSAMPEGYVQERTIFFKDDGNYKTRAEVKFEG
DTLVNRIELKGIDFKEDGNILGHKLEYNYNSHNVYIMADKQKNGIKVNFKIRHNIEDGSV
QLADHYQQNTPIGDGPVLLPDNHYLSTQSALSKDPNEKRDHMVLLEFVTAAGITLGMD
ELYKGLATLRESGPIYFETSLYKKAGTMTLILTSLLFFGLSLGPRTRVQAENLPKPILWAE
PGPVITWHNPVTIWCQGTLEAQGYRLDKEGNSMSRHILKTLESENKVKLSIPSMMWEHA
GRYHCYYQSPAGWSEPSDPLELVVTAYSRPTLSALPSPVVTSGVNVTLRCASRLGLGRF
TLIEEGDHRLSWTLNSHQHNHGKFQALFPMGPLTFSNRGTFRCYGYENNTPYVWSEPSD
PLQLLVSGVSRKPSLLTLQGPVVTPGENLTLQCGSDVGYIRYTLYKEGADGLPQRPGRQ
PQAGLSQANFTLSPVSRSYGGQYRCYGAHNVSSEWSAPSDPLDILIAGQISDRPSLSVQP
GPTVTSGEKVTLLCQSWDPMFTFLLTKEGAAHPPLRLRSMYGAHKYQAEFPMSPVTSA
HAGTYRCYGSRSSNPYLLSHPSEPLELVVSGATETLNPAQKKSDSKTAPHLQDYTVENLI
RMGVAGLVLLFLGILLFEAQHSQRSPPRCSQEANSRKDNAPFRVVEPWEQI*

*GPS-OSER1:*
MVSKGEELFTGVVPILVELDGDVNGHKFSVSGEGEGDATYGKLTLKFICTTGKLPVPWP
TLVTTLTYGVQCFSRYPDHMKQHDFFKSAMPEGYVQERTIFFKDDGNYKTRAEVKFEG
DTLVNRIELKGIDFKEDGNILGHKLEYNYNSHNVYIMADKQKNGIKVNFKIRHNIEDGSV
QLADHYQQNTPIGDGPVLLPDNHYLSTQSALSKDPNEKRDHMVLLEFVTAAGITLGMD
ELYKGLATLRESGPIYFETSLYKKAGTMKSEAKDGEEESLQTAFKKLRVDASGSVASLS
VGEGTGVRAPVRTATDDTKPKTTCASKDSWHGSTRKSSRGAVRTQRRRRSKSPVLHPP
KFIHCSTIASSSSSQLKHKSQTDSPDGSSGLGISSPKEFSAGESSTSLDANHTGAVVEPLRT
SVPRLPSESKKEDSSDATQVPQASLKASDLSDFQSVSKLNQGKPCTCIGKECQCKRWHD
MEVYSFSGLQSVPPLAPERRSTLEDYSQSLHARTLSGSPRSCSEQARVFVDDVTIEDLSG
YMEYYLYIPKKMSHMAEMMYT*

*GPS-ODC:*
MVSKGEELFTGVVPILVELDGDVNGHKFSVSGEGEGDATYGKLTLKFICTTGKLPVPWP
TLVTTLTYGVQCFSRYPDHMKQHDFFKSAMPEGYVQERTIFFKDDGNYKTRAEVKFEG
DTLVNRIELKGIDFKEDGNILGHKLEYNYNSHNVYIMADKQKNGIKVNFKIRHNIEDGSV
QLADHYQQNTPIGDGPVLLPDNHYLSTQSALSKDPNEKRDHMVLLEFVTAAGITLGMD
ELYKGLATLRESGPIYFETSLYKKAGTMNNFGNEEFDCHFLDEGFTAKDILDQKINEVSS
SDDKDAFYVADLGDILKKHLRWLKALPRVTPFYAVKCNDSKAIVKTLAATGTGFDCAS
KTEIQLVQSLGVPPERIIYANPCKQVSQIKYAANNGVQMMTFDSEVELMKVARAHPKA
KLVLRIATDDSKAVCRLSVKFGATLRTSRLLLERAKELNIDVVGVSFHVGSGCTDPETFV
QAISDARCVFDMGAEVGFSMYLLDIGGGFPGSEDVKLKFEEITGVINPALDKYFPSDSGV
RIIAEPGRYYVASAFTLAVNIIAKKIVLKEQTGSDDEDESSEQTFMYYVNDGVYGSFNCIL
YDHAHVKPLLQKRPKPDEKYYSSSIWGPTCDGLDRIVERCDLPEMHVGDWMLFENMG
AYTVAAASTFNGFQRPTIYYVMSGPAWQLMQQFQNPDFPPEVEEQDASTLPVSCAWES
GMKRHRAACASASINV*

*GPS-ODC K448X:*
MVSKGEELFTGVVPILVELDGDVNGHKFSVSGEGEGDATYGKLTLKFICTTGKLPVPWP
TLVTTLTYGVQCFSRYPDHMKQHDFFKSAMPEGYVQERTIFFKDDGNYKTRAEVKFEG

DTLVNRIELKGIDFKEDGNILGHKLEYNYNSHNVYIMADKQKNGIKVNFKIRHNIEDGSV
QLADHYQQNTPIGDGPVLLPDNHYLSTQSALSKDPNEKRDHMVLLEFVTAAGITLGMD
ELYKGLATLRESGPIYFETSLYKKAGTMNNFGNEEFDCHFLDEGFTAKDILDQKINEVSS
SDDKDAFYVADLGDILKKHLRWLKALPRVTPFYAVKCNDSKAIVKTLAATGTGFDCAS
KTEIQLVQSLGVPPERIIYANPCKQVSQIKYAANNGVQMMTFDSEVELMKVARAHPKA
KLVLRIATDDSKAVCRLSVKFGATLRTSRLLLERAKELNIDVVGVSFHVGSGCTDPETFV
QAISDARCVFDMGAEVGFSMYLLDIGGGFPGSEDVKLKFEEITGVINPALDKYFPSDSGV
RIIAEPGRYYVASAFTLAVNIIAKKIVLKEQTGSDDEDESSEQTFMYYVNDGVYGSFNCIL
YDHAHVKPLLQKRPKPDEKYYSSSIWGPTCDGLDRIVERCDLPEMHVGDWMLFENMG
AYTVAAASTFNGFQRPTIYYVMSGPAWQLMQQFQNPDFPPEVEEQDASTLPVSCAWES
GM*

*GPS-ODC W414C/415*:*
MVSKGEELFTGVVPILVELDGDVNGHKFSVSGEGEGDATYGKLTLKFICTTGKLPVPWP
TLVTTLTYGVQCFSRYPDHMKQHDFFKSAMPEGYVQERTIFFKDDGNYKTRAEVKFEG
DTLVNRIELKGIDFKEDGNILGHKLEYNYNSHNVYIMADKQKNGIKVNFKIRHNIEDGSV
QLADHYQQNTPIGDGPVLLPDNHYLSTQSALSKDPNEKRDHMVLLEFVTAAGITLGMD
ELYKGLATLRESGPIYFETSLYKKAGTMNNFGNEEFDCHFLDEGFTAKDILDQKINEVSS
SDDKDAFYVADLGDILKKHLRWLKALPRVTPFYAVKCNDSKAIVKTLAATGTGFDCAS
KTEIQLVQSLGVPPERIIYANPCKQVSQIKYAANNGVQMMTFDSEVELMKVARAHPKA
KLVLRIATDDSKAVCRLSVKFGATLRTSRLLLERAKELNIDVVGVSFHVGSGCTDPETFV
QAISDARCVFDMGAEVGFSMYLLDIGGGFPGSEDVKLKFEEITGVINPALDKYFPSDSGV
RIIAEPGRYYVASAFTLAVNIIAKKIVLKEQTGSDDEDESSEQTFMYYVNDGVYGSFNCIL
YDHAHVKPLLQKRPKPDEKYYSSSIWGPTCDGLDRIVERCDLPEMHVGDWMLFENMG
AYTVAAASTFNGFQRPTIYYVMSGPAC*

*GPS-ODC W414C/fs:*
MVSKGEELFTGVVPILVELDGDVNGHKFSVSGEGEGDATYGKLTLKFICTTGKLPVPWP
TLVTTLTYGVQCFSRYPDHMKQHDFFKSAMPEGYVQERTIFFKDDGNYKTRAEVKFEG
DTLVNRIELKGIDFKEDGNILGHKLEYNYNSHNVYIMADKQKNGIKVNFKIRHNIEDGSV
QLADHYQQNTPIGDGPVLLPDNHYLSTQSALSKDPNEKRDHMVLLEFVTAAGITLGMD
ELYKGLATLRESGPIYFETSLYKKAGTMNNFGNEEFDCHFLDEGFTAKDILDQKINEVSS
SDDKDAFYVADLGDILKKHLRWLKALPRVTPFYAVKCNDSKAIVKTLAATGTGFDCAS
KTEIQLVQSLGVPPERIIYANPCKQVSQIKYAANNGVQMMTFDSEVELMKVARAHPKA
KLVLRIATDDSKAVCRLSVKFGATLRTSRLLLERAKELNIDVVGVSFHVGSGCTDPETFV
QAISDARCVFDMGAEVGFSMYLLDIGGGFPGSEDVKLKFEEITGVINPALDKYFPSDSGV
RIIAEPGRYYVASAFTLAVNIIAKKIVLKEQTGSDDEDESSEQTFMYYVNDGVYGSFNCIL
YDHAHVKPLLQKRPKPDEKYYSSSIWGPTCDGLDRIVERCDLPEMHVGDWMLFENMG
AYTVAAASTFNGFQRPTIYYVMSGPACGNSCSNSRTPTSHPK*

*GPS-ODC Q419X:*
MVSKGEELFTGVVPILVELDGDVNGHKFSVSGEGEGDATYGKLTLKFICTTGKLPVPWP
TLVTTLTYGVQCFSRYPDHMKQHDFFKSAMPEGYVQERTIFFKDDGNYKTRAEVKFEG
DTLVNRIELKGIDFKEDGNILGHKLEYNYNSHNVYIMADKQKNGIKVNFKIRHNIEDGSV
QLADHYQQNTPIGDGPVLLPDNHYLSTQSALSKDPNEKRDHMVLLEFVTAAGITLGMD
ELYKGLATLRESGPIYFETSLYKKAGTMNNFGNEEFDCHFLDEGFTAKDILDQKINEVSS

SDDKDAFYVADLGDILKKHLRWLKALPRVTPFYAVKCNDSKAIVKTLAATGTGFDCAS
KTEIQLVQSLGVPPERIIYANPCKQVSQIKYAANNGVQMMTFDSEVELMKVARAHPKA
KLVLRIATDDSKAVCRLSVKFGATLRTSRLLLERAKELNIDVVGVSFHVGSGCTDPETFV
QAISDARCVFDMGAEVGFSMYLLDIGGGFPGSEDVKLKFEEITGVINPALDKYFPSDSGV
RIIAEPGRYYVASAFTLAVNIIAKKIVLKEQTGSDDEDESSEQTFMYYVNDGVYGSFNCIL
YDHAHVKPLLQKRPKPDEKYYSSSIWGPTCDGLDRIVERCDLPEMHVGDWMLFENMG
AYTVAAASTFNGFQRPTIYYVMSGPAWQLMQ*

*GPS-ODC W414X:*
MVSKGEELFTGVVPILVELDGDVNGHKFSVSGEGEGDATYGKLTLKFICTTGKLPVPWP
TLVTTLTYGVQCFSRYPDHMKQHDFFKSAMPEGYVQERTIFFKDDGNYKTRAEVKFEG
DTLVNRIELKGIDFKEDGNILGHKLEYNYNSHNVYIMADKQKNGIKVNFKIRHNIEDGSV
QLADHYQQNTPIGDGPVLLPDNHYLSTQSALSKDPNEKRDHMVLLEFVTAAGITLGMD
ELYKGLATLRESGPIYFETSLYKKAGTMNNFGNEEFDCHFLDEGFTAKDILDQKINEVSS
SDDKDAFYVADLGDILKKHLRWLKALPRVTPFYAVKCNDSKAIVKTLAATGTGFDCAS
KTEIQLVQSLGVPPERIIYANPCKQVSQIKYAANNGVQMMTFDSEVELMKVARAHPKA
KLVLRIATDDSKAVCRLSVKFGATLRTSRLLLERAKELNIDVVGVSFHVGSGCTDPETFV
QAISDARCVFDMGAEVGFSMYLLDIGGGFPGSEDVKLKFEEITGVINPALDKYFPSDSGV
RIIAEPGRYYVASAFTLAVNIIAKKIVLKEQTGSDDEDESSEQTFMYYVNDGVYGSFNCIL
YDHAHVKPLLQKRPKPDEKYYSSSIWGPTCDGLDRIVERCDLPEMHVGDWMLFENMG
AYTVAAASTFNGFQRPTIYYVMSGPA*

*3xFLAG-mTagBFP2-PSMD7-PSMD14:*
MDYKDHDGDYKDHDIDYKDDDDKTRMVSKGEELIKENMHMKLYMEGTVDNHHFKC
TSEGEGKPYEGTQTMRIKVVEGGPLPFAFDILATSFLYGSKTFINHTQGIPDFFKQSFPEGF
TWERVTTYEDGGVLTATQDTSLQDGCLIYNVKIRGVNFTSNGPVMQKKTLGWEAFTET
LYPADGGLEGRNDMALKLVGGSHLIANAKTTYRSKKPAKNLKMPGVYYVDYRLERIK
EANNETYVEQHEVAVARYCDLPSKLGHKLNTSMPELAVQKVVVHPLVLLSVVDHFNRI
GKVGNQKRVVGVLLGSWQKKVLDVSNSFAVPFDEDDKDDSVWFLDHDYLENMYGMF
KKVNARERIVGWYHTGPKLHKNDIAINELMKRYCPNSVLVIIDVKPKDLGLPTEAYISVE
EVHDDGTPTSKTFEHVTSEIGAEEAEEVGVEHLLRDIKDTGGGGSGGGGSGGGGSDTAE
QVYISSLALLKMLKHGRAGVPMEVMGLMLGEFVDDYTVRVIDVFAMPQSGTGVSVEA
VDPVFQAKMLDMLKQTGRPEMVVGWYHSHPGFGCWLSGVDINTQQSFEALSERAVAV
VVDPIQSVKGKVVIDAFRLINANMMVLGHEPRQTTSNLGHLNKPSIQALIHGLNRHYYSI
TINYRKNELEQKMLLNLH*

*SS-2xHA-LILRA4:*
MTLILTSLLFFGLSLGPRTRVQAENYPYDVPDYAGYPYDVPDYAGSYPYDVPDYALPKP
ILWAEPGPVITWHNPVTIWCQGTLEAQGYRLDKEGNSMSRHILKTLESENKVKLSIPSM
MWEHAGRYHCYYQSPAGWSEPSDPLELVVTAYSRPTLSALPSPVVTSGVNVTLRCASR
LGLGRFTLIEEGDHRLSWTLNSHQHNHGKFQALFPMGPLTFSNRGTFRCYGYENNTPYV
WSEPSDPLQLLVSGVSRKPSLLTLQGPVVTPGENLTLQCGSDVGYIRYTLYKEGADGLP
QRPGRQPQAGLSQANFTLSPVSRSYGGQYRCYGAHNVSSEWSAPSDPLDILIAGQISDRP
SLSVQPGPTVTSGEKVTLLCQSWDPMFTFLLTKEGAAHPPLRLRSMYGAHKYQAEFPM
SPVTSAHAGTYRCYGSRSSNPYLLSHPSEPLELVVSGATETLNPAQKKSDSKTAPHLQDY
TVENLIRMGVAGLVLLFLGILLFEAQHSQRSPPRCSQEANSRKDNAPFRVVEPWEQI*

Antibodies

We used the following antibodies: anti-GFP (Santa Cruz Biotech, sc-9996), anti-RFP/anti-tagBFP2 (Proteintech, pabr1), anti-cofilin (Cell Signaling Technology, 5175S), anti-PSMB5 (Thermo Fisher, A303-847A), anti-PSMC3 (Thermo Fisher, A303-538A), anti-vinculin (Cell Signaling Technology, 13901S), anti-FLAG (Sigma Aldrich, F1804), anti-alpha tubulin (Cell Signaling Technology, 2125S), and Alexa Fluor 647-conjugated anti-HA (Cell Signaling Technology, 3444S).

Lentiviruses and stable cell lines

To make lentiviruses, 6-well plates were transfected with plasmids encoding Gag, Pol, Rev, and VSV-G, along with the transfer plasmid using Polyjet (SignaGen). Media was changed after 24 hours, and the supernatant was harvested after 48 hours and frozen at -80 °C. Cells were transduced in 12-well plates with lentivirus along with 8 µg/mL polybrene (Millipore). After 48 hours, cells were split into media containing either 1 µg/mL puromycin (Thermo Fisher) or 50 µg/mL hygromycin B (Thermo Fisher). Selection typically required ~5 days. For validating E3 ligases, the GPS reporters were first introduced and selected, followed by guides.

Lentivirus for screens were packaged in 15-cm plates using Polyjet (SignaGen). Media was changed after 24 hours, and supernatant was harvested at both 48 hours and 72 hours, pooled, concentrated using Lenti-X Concentrator (Clontech), and frozen at -80 °C. Multiplicity of infection (MOI) was calculated by transducing cells with various amounts of concentrated lentivirus. The MOI for GPS libraries were calculated as the fraction of cells that were positive for DsRED. The MOI for CRISPR libraries were calculated by growing cells either with or without 1 µg/mL puromycin. After cells were fully selected (typically 3-4 days), the MOI was calculated as the ratio between the number of cells grown with puromycin divided by the number of cells grown without puromycin.

Pooled GPS screening

*GPS library construction*

For the cysteine-wide GPS library, a pool of 130,475 oligos was synthesized by Agilent Technologies. The sequences are derived from 20,593 unique cysteine motifs from the human proteome: 6,923 CxC motifs and 13,706 CxxC motifs. Each motif is present in the library in three different registers, each of 34 amino acids but shifted by 5 amino acids relative to one another. Every wild-type peptide (e.g. CxC or CxxC) also had a paired mutant version (e.g. AxC or AxxC). We identified 7,379 cysteine peptides within our library that are derived from C2H2 zinc fingers and also included paired mutants that disrupted the histidines (i.e. CCAA). For the site-saturating mutagenesis library, approximately 66 peptides were selected based on their cysteine-dependent degradation ($PSI^{Ala}$ - $PSI^{Cys}$), as well as whether other registers of the same peptide scored highly. A library of all single-substitution mutations was synthesized by Agilent and cloned into GPS3.0-I24-Puro.

Nearly all of the site-saturating mutagenesis was performed on the original peptide sequences from the cysteine-dependence GPS screen. Note, however, that we had first identified ATE1_1 in Timms et al., 2023 as a cysteine-dependent substrate (*18*). Therefore, in the saturation mutagenesis and CRISPR knockout screens, we included the constant flanking sequence (underlined) that we had used in the prior study:

DYFPSEDFYR**CGYC**KNESGSRSNGRIARAKASTN*. This differed from the top scoring ATE1 peptide that was in our original GPS library:
SVVDYFPSEDFYR**CGYC**KNESGSRSNGMWAHSMT*.
Libraries were amplified using Q5 DNA Polymerase (NEB), cloned into GPS3.0-I24-Puro using Hifi DNA Assembly (NEB), concentrated using AMPure beads (Beckman Coulter), and electroporated into ElectroMAX DH10B cells (Invitrogen) while maintaining 500X representation.

*Cell culture and sorting*

Each screen replicate was performed using the same lentiviral library, but was independently transduced, cultured, and sorted, maintaining a minimum representation of 400X per library member throughout. For each replicate, twenty 15-cm plates were seeded with 8 million cells per plate one day prior to transduction. For the cysteine-wide GPS screen, we used independent wild-type HEK-293T clones, as well as two APPBP2 knockout clones (KO3 and KO11). Other screens were performed with wild-type HEK-293T.

On the day of transduction, cells were incubated with 8 µg/mL polybrene and lentivirus at a multiplicity of infection (MOI) of ~0.2. Two days post-transduction, cells were selected in 1.5 µg/mL puromycin for approximately 7 days, with passaging every 2-3 days. Cells were then transferred to non-selective media for 1-2 days before sorting. For the pharmacological screen, cells were split into separate arms that were treated prior to sorting with DMSO, 1 µM bortezomib (8 h), 1 µM TAK-243 (8 h), or 1 µM MLN-4924 (12 h).

Cells were harvested for flow cytometry 10 or 11 days post-transduction. Cells were sorted on a MoFlo Astrios Cell Sorter (Beckman Coulter) based on the GFP:DsRed fluorescence ratio. Six evenly spaced bins were defined, and approximately 10 million cells were typically collected per bin. Sorted cells were pelleted and stored at −80 °C prior to genomic DNA extraction. Bins with low cell numbers were supplemented with wild-type 293T cells after sorting to facilitate efficient genomic isolation.

Genomic DNA was isolated using the Gentra Puregene Cell Kit (Qiagen), sequences were amplified using Q5 High-Fidelity DNA Polymerase (NEB), and 150bp Illumina sequencing was performed at the Harvard Medical School Biopolymers Facility.

*Analysis of GPS results*

Sequencing reads were trimmed using cutadapt (*67*), and aligned using bowtie2 (*68*). We filtered for peptides with at least 100 reads across all bins and normalized read counts to be proportional to the number of cells collected from each sorting bin. Protein stability index (PSI) scores were then calculated as the average bin number for a given peptide.
Since the stability of the library as a whole was dramatically affected by bortezomib and TAK-243 treatment, we used a custom script to adjust the "cell counts" in order to minimize the overall differences in PSI scores between TAK-243 and bortezomib, and between MLN-4924 and DMSO. Ubiquitin-independent substrates were defined as $PSI^{BORT}$ - $PSI^{TAK} \geq 0.5$, while Cullin-dependent substrates were defined as $PSI^{MLN}$ - $PSI^{DMSO} \geq 0.5$.

CRISPR knockout screening

The genome-wide CRISPR knockout library was a gift of David Root and was previously used (*13*). The ubiquitin-wide CRISPR knockout library was also previously described (*69*).

CRISPR screening was performed similarly to the GPS stability screens, with the following modifications. We transduced cells with an MOI of 0.3, maintaining at least 1000X representation throughout. Cells were harvested 7-10 days post-transduction and sorted on a MA-900 (Sony). The highest 5% of cells by GFP:DsRED fluorescent ratio were collected as the enriched population and an unsorted population was also collected. Sequencing was performed on either an Illumina or AVITI platform (Element Biosciences). Reads were trimmed and aligned to the sgRNA reference library, and gene-level enrichment scores were calculated using MAGeCK (*70*).

Base editor CRISPR screening

*Base editor library construction*

We downloaded from UCSC genome browser the hg38 genomic DNA sequences for each of the 19S genes, including PSMD14, along with USP14, UCHL5, MIDN, TXNL1. We determined all 20nt guide sequences that overlapped with the CDS region and had an NG PAM. The reporter sequence was created by appending the 6bp flanking sequence to both sides of the 20nt guide sequence. Guides that started with a G had this first nucleotide removed. Splice-targeting guides (positive controls) were determined as containing a splice site within the 32nt reporter sequence. We also designed 727 negative control guides that collectively targeted 20 different intergenic or intronic regions. A subpool of guides targeting PSMD14, tiling every nucleotide was also created along with separate negative control guides.

DNA oligonucleotides containing the guide, reporter, and barcode were synthesized by Twist Bioscience and cloned into pLentiCRISPR-ABE8e-dCas9SpRY or pLentiCRISPR-ABE8e-dCas9NG as previously described (*35*), and lentivirus was packaged as described above.

*Cell culture and sorting*

The NG and SpRY libraries were pooled at a ratio of 85%:15%. HEK-293T cells were seeded into four 15-cm plates the day prior to transduction at 6.25 million cells per plate. The following day, concentrated lentivirus and polybrene were added, resulting in an MOI of 0.5, and maintaining 1000X representation throughout all steps. After 24 hours the media was replaced with fresh media containing 2 mM valproic acid (VPA). After another 48 hours, cells were split into media containing 1 ug/mL puromycin. Cells were maintained in selection until the day prior to sorting, with sorting occurring 9 days following transduction. The highest 1% of cells by GFP:DsRED fluorescent ratio were collected as the enriched population along with an unsorted population.

*Base editor analysis*

Sequencing was performed on the AVITI platform (Element Biosciences) by Quintara Biosciences (Cambridge, MA). Reads were trimmed and aligned to either the NG reference library, allowing for no mismatches, or the SpRY reference library, allowing for up to 2bp substitutions to account for possible editing of the guide sequence.

Editing outcomes for each of the guides were determined using the crispr-bean algorithm (*35*), with mutations having to be present in at least 5% of alleles and 20% of samples. Enrichment scores for each guide was determined by first filtering for those that had at least 200 reads in the inputs of each of the replicate screens. Then, enrichment scores were calculated by normalizing the total number of reads in each population and then dividing the read counts in the sorted population by the read counts in the input population. Per-residue enrichment scores for each residue in the 19S proteasome were calculated by multiplying one minus the enrichment score of a guide by the frequency of each particular residue being mutated, summed across all guides.

Individual GPS reporter assays

Individual validation assays were performed in 6-well or 12-well plates. HEK-293T cells stably expressing GPS reporters were detached using 200 uL trypsin-EDTA, quenched using 200 uL growth medium, and measured with a Cytoflex LX (Beckman Coulter) on the FITC and PE channels. Analysis was performed on Flowjo 10.10. Cells were gated on singlets, followed by DsRED positive cells. The GFP to DsRED ratios were then calculated and presented as a histogram.

Cells were treated with drugs for 6-8 hours prior to flow cytometry. The following concentrations were used: 1 μM bortezomib (APExBIO), 1 μM TAK-243 (Selleckchem), 1 μM MLN-4924 (Selleckchem), and 5 μM N,N,N′,N′-tetrakis(2-pyridinylmethyl)-1,2-ethanediamine (TPEN, Sigma). Drugs were added concurrently in cases where more than one was used.

To stain for membrane-localized SS-2xHA-LILRA4, trypsonized HEK-293T cells were incubated with anti-HA antibody conjugated to Alexa Fluor 647 (Cell Signaling Technology) diluted 1:100 in PBS at 4 °C for 30 minutes, washed, and then analyzed on a Cytoflex LX (Beckman Coulter).

CRISPR knockouts (E3 ligase validation)

The following guides were annealed and ligated into pLentiCRISPRv2, and lentivirus encoding Cas9 and guides were produced.

sgAAVS1: GGGGCCACTAGGGACAGGAT
sgFBXO3_1: GATGCTGTGGAAGCGCAGAT
sgFBXO3:_2: GTATCAAGTCATGATCCGCTG
sgFBXO21_1: GTCTCTGGCCAGATTCCCAGG
sgFBXO21_2: GCTTGTTTGCAAAACCCTTCG
sgAMBRA1_1: GTGGTAGAAGATAAAACCCGG
sgAMBRA1_2: GCCATAATATCTATATTACGG
sgTRPC4AP_1: GTAACATTCTGCTGCAGCTG
sgTRPC4AP_2: GACACTCAACAAAGTCACTG
sgPPIL2_1: GCAGGGGACACTCACCTGGA
sgPPIL2_2: GCACCAACAACACCCACATCG

Cell lines carrying GPS reporters were transduced with lentivirus, cells were selected using puromycin, and protein stability was assessed by flow cytometry on a Cytoflex LX (Beckman Coulter).

Purification of recombinant GFP substrate

We purified recombinant GFP fusions from bacteria using the same protein sequences as in the cellular GPS reporters, except with the addition of a 1xFLAG tag to the N-terminus of GFP. Protein was purified using the Impact system (NEB), according to the manufacturer's protocol. Briefly, BL21(DE3) cells were transformed with pTXB1 constructs carrying the various GFP fusion, grown at 37 °C, and induced with 0.2 mM IPTG. Bacteria were lysed with B-PER supplemented with 2U/mL benzonase, 1 mM PMSF, and 5 mM EDTA. After clarification at 30,000 xg, cleared lysate was incubated with chitin resin and washed thoroughly with 500 mM NaCl, 25 mM Hepes 8.5, and 5 mM EDTA. GFP fusion protein was eluted with 50 mM DTT overnight at RT, frozen in liquid nitrogen, and stored at -80 °C.

PSMD7-PSMD14 pulldown assay

HEK 293T cells were seeded in 15-cm plates and transfected with 30 μg of pMAX 3xFLAG-tagBFP2-PSMD7(1-182)-PSMD14(26-221) fusion construct DNA per plate using 90 μL of PolyJet (SignaGen), following the manufacturer's protocol. After 48 hours, cells were harvested and lysed in 1.7 mL of lysis buffer per plate (100 mM NaCl, 50 mM HEPES (pH 7.5), 1 μM bortezomib, 0.1% Triton X-100, 1 mM DTT, and Pierce Complete EDTA-free protease inhibitor). Lysates were incubated with rotation at 4 °C for 10 minutes and clarified by centrifugation at 21,000 × g for 30 minutes at 4 °C.

To reduce nonspecific binding, 600 μL of clarified lysate was precleared by incubating with 50 μL (slurry volume) of control magnetic agarose beads (Chromotek) and 5% (w/v) bovine serum albumin (BSA) for 1 hour at 4 °C.

In parallel, 25 μL slurry volume of GFP-Trap magnetic beads were incubated in ysis buffer supplemented with 5% (w/v) BSA and 120 μg of recombinant EGFP fusion protein for 30 minutes at 4 °C. Unbound EGFP was removed by washing, and then precleared PSMD7-PSMD14-containing lysates were incubated with EGFP-loaded GFP-Trap beads for 30 minutes at 4 °C with rotation. Beads were washed three times with lysis buffer, transferred to new microcentrifuge tubes, and eluted by boiling in NuPAGE LDS Sample Buffer (Thermo Fisher) supplemented with 50 mM DTT.

Eluted proteins were separated using NuPAGE Bis-Tris gels (Thermo Fisher), transferred to nitrocellulose membranes, and detected by chemiluminescence.

Cycloheximide chase assay

Cell lines stably expressing GPS reporters were seeded into 12-well plates and cultured until confluent. Cells were then treated with 100 μg/mL cycloheximide alone, or in combination with either 1 μM bortezomib or 1 μM TAK-243, and harvested at indicated timepoints.

At each timepoint, cells were rinsed once with PBS and lysed directly in-well with 200 μL of NuPAGE LDS Sample Buffer (Thermo Fisher) supplemented with 50 mM DTT. Lysates were denatured at 100 °C and subjected to probe sonication to shear genomic DNA. Proteins were resolved by SDS-PAGE using NuPAGE Bis-Tris gels (Thermo Fisher), transferred to nitrocellulose membranes via wet transfer, and detected by chemiluminescence.

Generation of APPBP2 knockout cell lines

HEK-293T cells were transfected with plasmids encoding GFP, Cas9, and dual guides. These guides were designed to excise either a region of exon 1 including the start codon, or separately targeting exon 2, resulting in a frameshift if removed. Cells were single-cell sorted into 96-well plates and individual clones were expanded. Knockouts were determined by PCR of genomic regions, as well as by stabilization of a model RxxG substrate by GPS.

Sample preparation LFQ quantitative mass spectrometry

Cells were treated with 250 nM Bortezomib, 1 μM MLN7243, 0.5 μM MLN4924, or DMSO vehicle control for 6 hours. Treated cells were harvested and washed twice with PBS before snap freezing in liquid nitrogen.

Cells were lysed by addition of lysis buffer (8 M Urea, 50 mM NaCl, 50 mM 4-(2-hydroxyethyl)-1-piperazineethanesulfonic acid (EPPS) pH 8.5, Protease and Phosphatase inhibitors) and homogenization by bead beating (BioSpec) for three repeats of 30 seconds at 2400 strokes/min. Bradford assay was used to determine the final protein concentration in the clarified cell lysate. Fifty micrograms of protein for each sample was reduced, alkylated and precipitated using methanol/chloroform as previously described (*71*), and the resulting washed precipitated protein was allowed to air dry. Precipitated protein was resuspended in 4 M urea, 50 mM HEPES pH 7.4, followed by dilution to 1 M urea with the addition of 200 mM EPPS, pH 8. Proteins were digested with the addition of LysC (1:50; enzyme:protein) and trypsin (1:50; enzyme:protein) for 12 h at 37 °C. Sample digests were acidified with formic acid to a pH of 2-3 before desalting using C18 solid phase extraction plates (SOLA, Thermo Fisher Scientific). Desalted peptides were dried in a vacuum-centrifuged and reconstituted in 0.1% formic acid for liquid chromatography-mass spectrometry analysis.

Data were collected using a TimsTOF Ultra (Bruker Daltonics, Bremen, Germany) coupled to a nanoElute LC pump (Bruker Daltonics, Bremen, Germany) via a CaptiveSpray nano-electrospray source. Peptides were separated on a reversed-phase C18 column (25 cm x 75 µm ID, 1.6 µM, IonOpticks, Australia) containing an integrated captive spray emitter. Peptides were separated using a 50 min gradient of 2 - 30% buffer B (acetonitrile in 0.1% formic acid) with a flow rate of 250 nL/min and column temperature maintained at 50 ºC.
The TIMS elution voltages were calibrated linearly with three points (Agilent ESI-L Tuning Mix Ions; 622, 922, 1,222 m/z) to determine the reduced ion mobility coefficients (1/K0). To perform diaPASEF, we used py_diAID (*72*), a python package, to assess the precursor distribution in the m/z-ion mobility plane to generate a diaPASEF acquisition scheme with variable window isolation widths that are aligned to the precursor density in m/z. Data was acquired using twenty cycles with three mobility window scans each (creating 60 windows) covering the diagonal scan line for doubly and triply charged precursors, with singly charged precursors able to be excluded by their position in the m/z-ion mobility plane. These precursor isolation windows were defined between 350 - 1250 m/z and 1/k0 of 0.6 - 1.45 V.s/cm2.

<u>LC-MS data analysis</u>

The diaPASEF raw file processing and controlling peptide and protein level false discovery rates, assembling proteins from peptides, and protein quantification from peptides were performed in library free mode searched against a Swissprot human database (January 2021). Database search criteria largely followed the default settings for directDIA including: tryptic with two missed cleavages, carbamidomethylation of cysteine, and oxidation of methionine and precursor Q-value (FDR) cut-off of 0.01. Precursor quantification strategy was set to Robust LC (high accuracy) with RT-dependent cross run normalization. Proteins with low sum of abundance (<2,000 x no. of treatments) were excluded from further analysis and resulting data was filtered to only include proteins that had a minimum of 3 counts in at least 3 replicates of each independent comparison of treatment sample to the DMSO control. Proteins with missing values were imputed by random selection from a Gaussian distribution either with a mean of the non-missing values for that treatment group or with a mean equal to the median of the background (in cases when all values for a treatment group are missing) (*73*). Protein abundances were scaled and significant changes compared to the DMSO control were assessed by two-sided moderated t-test as implemented in the limma package within the R framework (*74*).

Purification of PSMD7-PSMD14

His-tagged-GST-PSMD7 and PSMD14 were coexpressed in E. coli after growing to OD600~0.6 in dYT and inducing with 0.5 mM IPTG overnight at 18 degrees C. Cells were lysed by sonication, clarified, and purified by Ni-affinity chromatography using Ni-NTA (ThermoFisher) using Ni A Buffer (50 mM HEPES 7.6, 200 mM NaCl, 20 mM imidazole), and Ni B Buffer (Ni A+ 250 mM imidazole). Fractions containing the heterodimer were concentrated, and passed through a Superdex S200 16/600 (Cytiva) column, concentrated and the concentration determined by the extinction coefficient.

Inhibition of ScRpn11 deubiquitinase activity

*S. cerevisiae* Rpn11 deubiquitinase activity was measured using Ubiquitin- 5-carboxytetramethylrhodamine (Ub-TAMRA) as a substrate as previously described (*38*, *39*). Briefly, *Sc* Rpn8/Rpn11 heterodimer and the Ubiquitin-GGGC substrate was expressed and purified from E. coli and the Ubiquitin substrate was attached to a TAMRA maleimide using cysteine chemistry. The fluorescence polarization (FP) of the TAMRA dye as it is liberated from Ub-TAMRA was monitored in a 384 well plate (Corning 3844) using a Clariostar Plus (BMG Labtech) with $\lambda_{excitation}$ = 540 nm (band pass 20 nm) and $\lambda_{emission}$= 590 nm (band pass 20nm). All proteins were purified and assayed in GF buffer (50 mM HEPES 7.6, 50 mM NaCl, 50 mM KCl, 10 mM MgCl2, 5% glycerol). 0.5 uM *Sc* Rpn8/11 was added to 100 uM Ub-TAMRA with and without 100 uM peptide. The slope of FP loss was calculated and an $IC_{50}$ was calculated using Prism (Graphpad) fitting to: Y=Bottom+(Top-Bottom)/(1+(X/IC50))).

Purification of S. cerevisiae 26S proteasome

Proteasome was purified as previously described using M2 anti-FLAG affinity resin (Sigma) (*75*). Briefly, a strain of yeast containing a N-terminal FLAG on Rpn11 (YYS40) was grown in YPD media, frozen, lysed via cryo-milling and then clarified by spinning at 26,000 xg for 30 minutes at 4° C. The supernatant was applied to anti-FLAG resin, washed in buffer A (60mM HEPES pH 7.6, 25 mM NaCl, 25 mM KCl, 5 mM MgCl2, 0.5 mM EDTA, 5% glycerol and 2 mM ATP) and proteasome was eluted by the addition of Buffer A plus 0.15 mg/mL FLAG peptide (Genscript). Proteasome was then applied to a Superose 6 increase 10/300 (Cytiva) column equilibrated in GF buffer supplemented with 0.5 mM ATP. Protein concentration was determined using Bradford (Bio-Rad).

Measurement of 26S ATPase rates

ATP-hydrolysis rates of 26S proteasome were determined using an NADH-coupled assay as described previously (*76*, *77*). 100 nM 26S proteasome was added to 100 uM peptide and a 1X ATPase mix (final concentrations: 1 mM NADH, 2.5 mM ATP, 7.5 mM phosphoenolpyruvate, 3 U/mL pyruvate kinase, and 3 U/mL lactate dehydrogenase). NADH consumption was measured in a clear 384 well plate (Corning 3544) by measuring the absorbance at 340 nm in a Synergy Neo2 Multi-Mode Plate Reader (Biotek).

Additional Analysis

In vitro bar plots and titration curves were generated using Graphpad Prism (version 10.3.1). All other plots were generated using the ggplot2 package (version 3.5.2) within R (version 4.4.3). Percent identity between human cellular FOS and Finkel-Biskis-Jinkins murine osteosarcoma virus FOS were determined using the EMBOSS Needle Pairwise Sequence Alignment from the EMBL. ChatGPT (OpenAI) was used to improve writing clarity; all changes were reviewed for accuracy.

#### Supplementary Text

Specificity in targeting double cysteine motifs

FBXO3, FBXO21, and the FEM1A/B/C family each recognized relatively specific degrons composed of the CXXC motifs in combination with at least two other critical residues (Fig. 2B, fig. S3A). For instance, the two substrates of FBXO21 both contain a CGYC motif, and the FBXO3 substrate is recognized through a LCxGCxEC motif. By contrast, AMBRA1 and TRPC4AP primarily recognized just the two cysteine residues. Consistent with their ability to broadly target CXXC-containing sequences, AMBRA1 and TRPC4AP collectively scored in 8 of the 11 screens that we performed (Fig. 2D, fig. S3B). In several cases they even appeared as secondary hits where another E3 was more dominant, suggesting that they may act redundantly or serve as a backup surveillance mechanism.

Notably, in all cases where TRPC4AP was identified as a hit, the U-box E3 ligase PPIL2 was also a hit and was required for efficient degradation (Fig. 2D; fig. S3B, D), suggesting that TRPC4AP and PPIL2 may function together to ubiquitylate substrates in the same pathway. One possibility is that PPIL2 may work to extend the ubiquitin chains that TRPC4AP initiates, allowing for efficient proteasomal targeting; indeed, the stabilization of TRPC4AP substrates by knocking out PPIL2 was more modest than for knocking out TRPC4AP, possibly suggesting an accessory role for PPIL2 in degrading cysteine-containing TRPC4AP substrates.

Role of APPBP2 in targeting cysteines

We investigated substrates of APPBP2, an E3 ligase previously implicated in cysteine-dependent degradation (*18*). APPBP2 is known to recognize C-terminal degrons (C-degrons) ending in RXXG motifs (*66*), but how it recognizes internal cysteine motifs is unclear. We identified and characterized two new substrates from our screens that were degraded in a cysteine-dependent manner and required APPBP2 for degradation: NFXL1-15 (fig. S3A-C) and TG-2 (fig. S3D-F). Our bin-level analysis of the screening data confirmed that they were degraded in a cysteine- and APPBP2-dependent manner (fig. S3B, E), and that they were degraded in a manner dependent on Cullin-RING ligases, ubiquitin conjugation, and the proteasome (fig. S3C, F). Notably, site-saturating mutagenesis indicated that both NFXL1-15 and TG-2 depended on not just a CxC motif, but were also dependent on sequences in the very C-terminal 5-8 amino acids. These C-terminal terminal sequences were not the canonical RXXG motif, but rather were a complex sequence motif rich in arginine, proline, and histidine residues.

We also investigated in detail four APPBP2 substrates that we identified previously: GMCL1(6), MATN2(64), MLL5(27), and TRIM35(10) (*18*). We validated that they were degraded in a cysteine-dependent manner by APPBP2 (fig. S4G-I). Using GMCL1(6) as a model substrate, we performed site-saturating mutagenesis and noted that it, too, depended on both an internal CxC motif and a constant C-terminal motif (fig. S4G). This C-terminal motif was the constant sequence (-RIARAKASTN*) added to all of the peptides in our prior study (*18*). As with the other APPBP2 substrates, neither the CxC or the C-terminal motif alone was sufficient for degradation (fig. S4G). We also removed the C-terminal constant sequence for each of these four substrates, verifying that degradation by APPBP2 depends on this terminal sequence (fig. S4J).

Collectively, this data suggests that neither motif – the cysteine motif nor the C-terminal motif – is alone sufficient for degradation; instead, a bipartite degron is necessary. This may also explain why we observed few peptides in our cysteine-wide GPS library (Fig. 1) that were degraded by APPBP2 in a cysteine-dependent manner. This may likely reflect that APPBP2 requires specific C-terminal sequences in conjunction with the internal cysteines. Since APPBP2 can require 5-8 specific amino acids at the C-terminus, presumably few peptides in our cysteine-wide GPS library have such a sequence in combination with the right internal cysteine motif. By contrast, the C-terminal constant sequence that was used in Timms et al., 2023 may have some basal affinity for APPBP2, resulting in a higher frequency of APPBP2 substrates.

We next wanted to investigate how APPBP2 recognizes these bipartite cysteine substrates. Through AlphaFold modeling we found that APPBP2 has two residues, Cys305 and His343, near its canonical C-degron recognition site that may bind the paired cysteine substrates through a coordinated zinc (fig. S4K). Indeed, mutating the His343 residue blocked degradation of the cysteine-dependent substrate, but did not affect the degradation of a canonical RXXG substrate that does not use this site (fig. S4L). Addition of the cell-permeable chelator TPEN likewise blocked degradation of the cysteine-containing substrate MATN2(64) (fig. S4M). We also performed co-immunoprecipitation experiments and found that the addition of the metal chelators EDTA or ortho-phenanthroline (OPT) to lysate blocked the interaction of GMCL1(6) to APPBP2, as did mutation of the APPBP2 residue His343 (fig. S4N). Importantly, we found that these treatments did not affect the interaction with a canonical RXXG substrate. Together, we conclude that APPBP2 contains a zinc-binding site that can recruit substrates that possess double cysteine motifs in conjunction with specific C-terminal sequences.

Redundancy between pathways

While most cysteine degrons adhered to either Cullin-dependent or ubiquitin-independent pathways (fig. S6C), we also uncovered cases where slight changes in sequence could alter the degradation pathway of a substrate. For example, homologous peptides from GATA3, GATA4, and GATA6 showed varying dependence on the two pathways, with GATA3 being predominantly Cullin-dependent and GATA6 being predominantly ubiquitin-independent (fig. S6I-N).

While certain degron features are characteristic of each pathway (e.g. nearby arginines, serines, and threonines for ubiquitin-independent degradation), mutations outside of the degron may lead to subtle effects – including to cysteine reactivity, exposure, and secondary structure – that may affect its degradation behavior. Additional work will be needed to accurately predict the protein stability and pathway dependence of a given sequence.

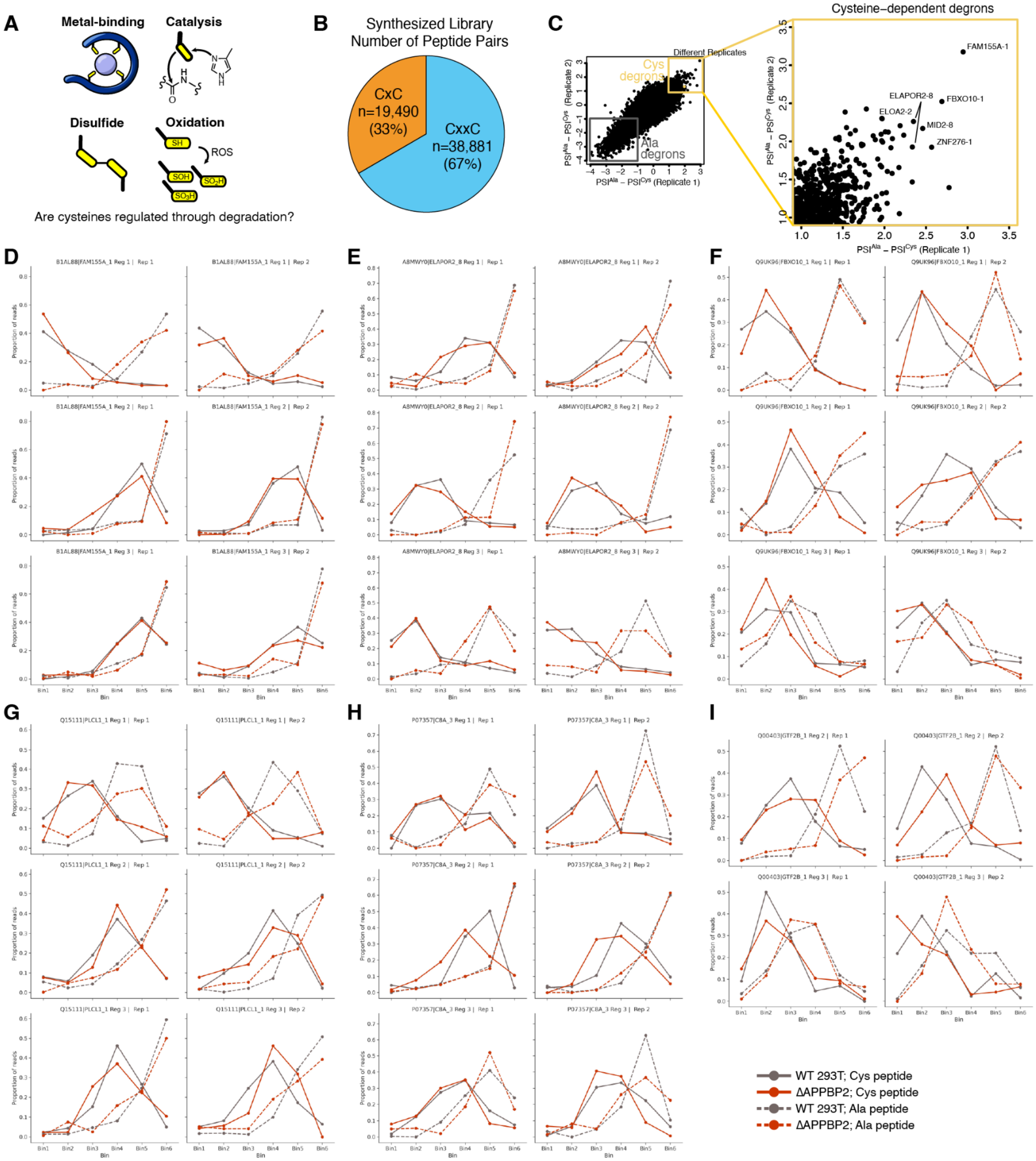


### Supplementary Figure 1

**Fig. S1. Library design and screening of cysteine-dependent degrons**. (**A**) Cysteine is among the most chemically versatile amino acids: it binds to metals, participates in catalysis, forms disulfide bonds and other covalent crosslinks, and is readily oxidized. (**B**) The peptide library includes both CxC and CxxC motifs derived from the human proteome. (**C**) Cysteine-dependence degradation scores ($PSI^{Ala}$ - $PSI^{Cys}$) from two biological replicates are plotted against each other. The inset highlights cysteine-dependent peptides that score greater than 1.0 in both replicates. (**D**-**I**) Representative cysteine-dependent hits are shown with relative read distributions across the six flow cytometry bins. Each peptide was tested in two biological replicates (Rep 1 and Rep 2) and across three overlapping sequence windows (Reg 1, Reg 2, and Reg 3). Solid lines represent wild-type peptides; dashed lines show cysteine-to-alanine mutants. Screens were performed in wild-type HEK293T cells (gray lines) and APPBP2 knockout cells (red lines).

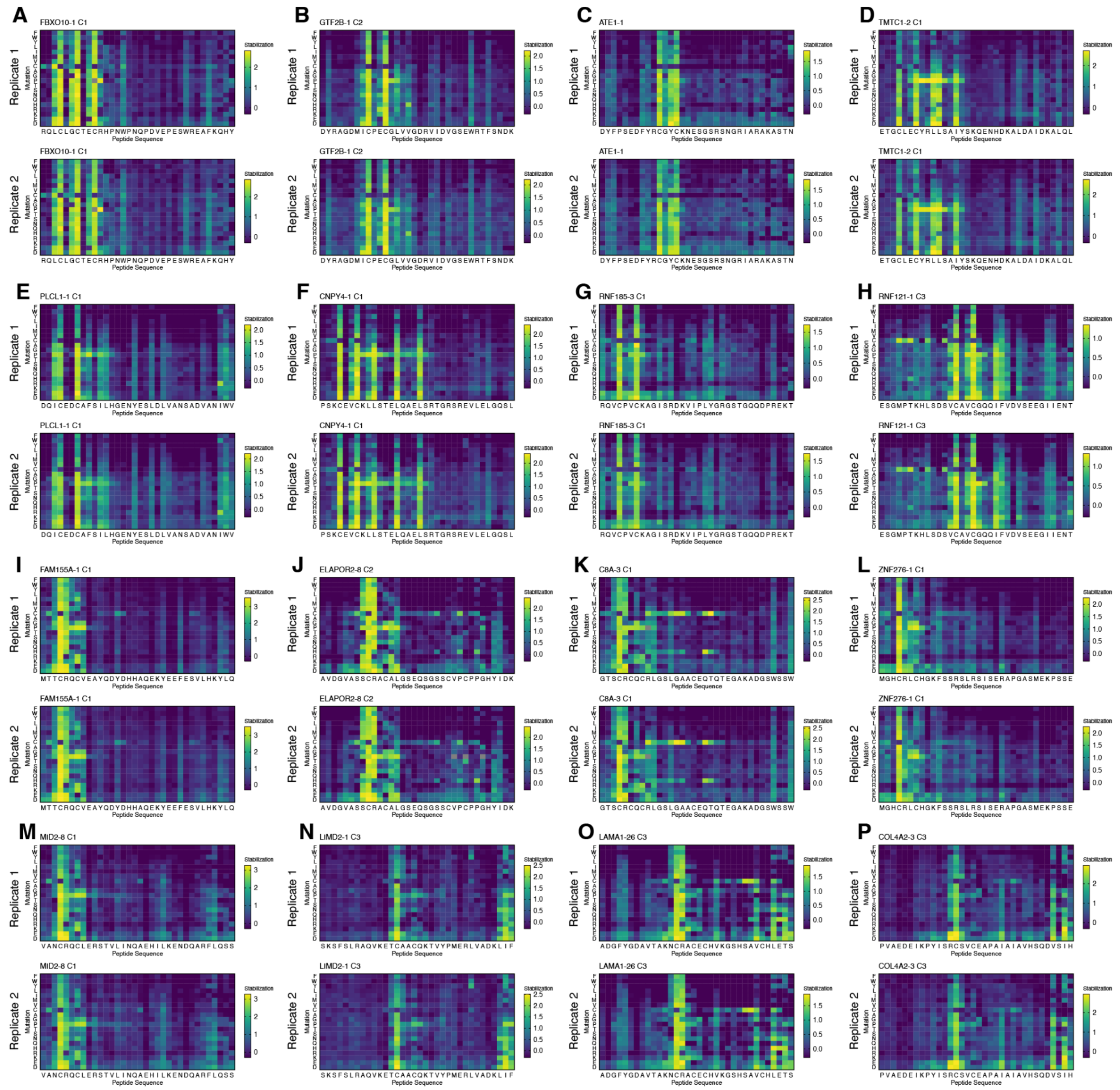

### Supplementary Figure 2

**Fig. S2. Additional saturation mutagenesis heatmaps. (A–H)** Heatmaps for degrons requiring two or more cysteines. **(I–P)** Heatmaps for degrons dependent on a single cysteine residue. In each heatmap, the original peptide sequence is shown along the X-axis, and the amino acid substitutions are on the Y-axis. Yellow indicates substitutions that stabilize the reporter. Biological replicates exhibited high concordance.

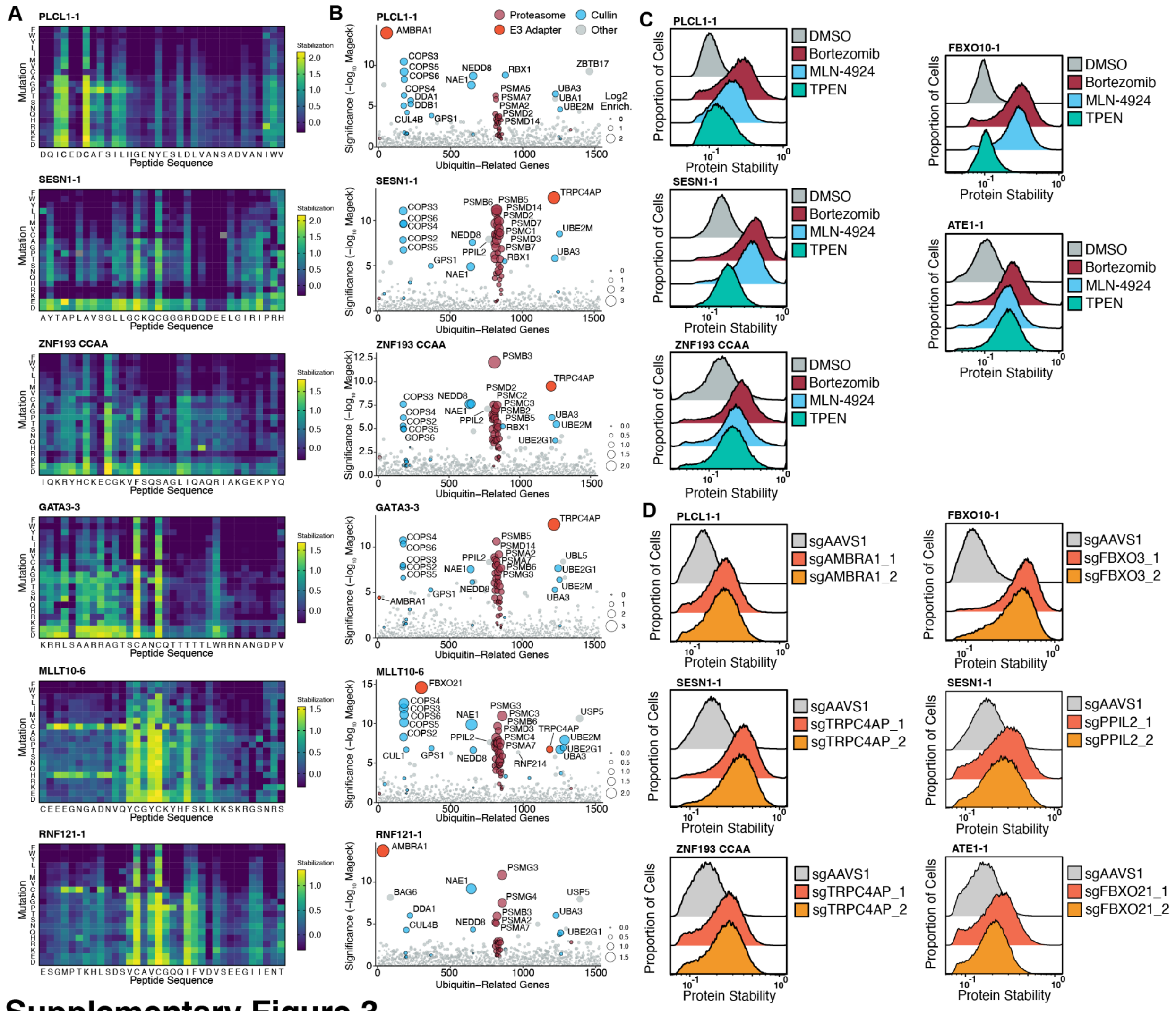


#### Supplementary Figure 3

**Fig. S3. Cullin-RING E3 ligases recognize a variety of paired cysteine substrates.** (**A**) Site-saturation mutagenesis heatmaps for a panel of substrates containing double cysteine degrons. (**B**) CRISPR-Cas9 screen results using a ubiquitin-focused library, showing specific E3 ligases required for degradation of each substrate. (**C**) Individual peptide stability assays were performed in the presence of bortezomib , MLN-4924 , or TPEN. Tight binding to zinc through the three cysteines of FBXO10-1 may explain its resistance to TPEN. (**D**) Polyclonal CRISPR-Cas9 knockout of individual E3 ligases in cells expressing stability reporters.

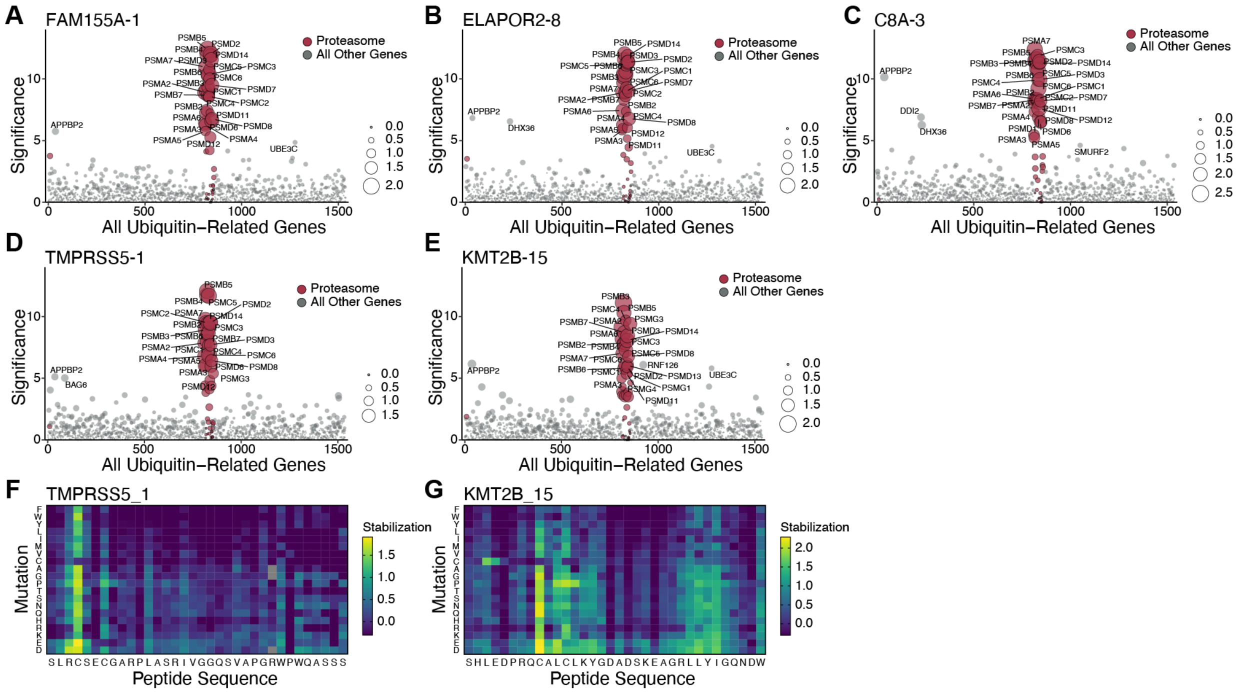


### Supplementary Figure 4

**Fig. S4. Ubiquitin-focused CRISPR-Cas9 screens of single cysteine substrates.** (**A-E**) CRISPR-Cas9 knockout screens were performed using a focused ubiquitin pathway library on five distinct substrates bearing single-cysteine degrons. Each dot represents a gene, dot size indicates sgRNA enrichment, and statistical significance (−$\log_{10}$[MAGeCK positive score]) is shown on the Y-axis. Genes encoding proteasomal subunits are highlighted in red. APPBP2 frequently scores in our screens as a non-specific hit; these substrates are not degraded via APPBP2 (see also fig. S1). (**F**, **G**) Site-saturation mutagenesis heatmap of TRPRSS5-1 (F) and KMT2B-15 (G), showing that they are degraded through a single cysteine motif.

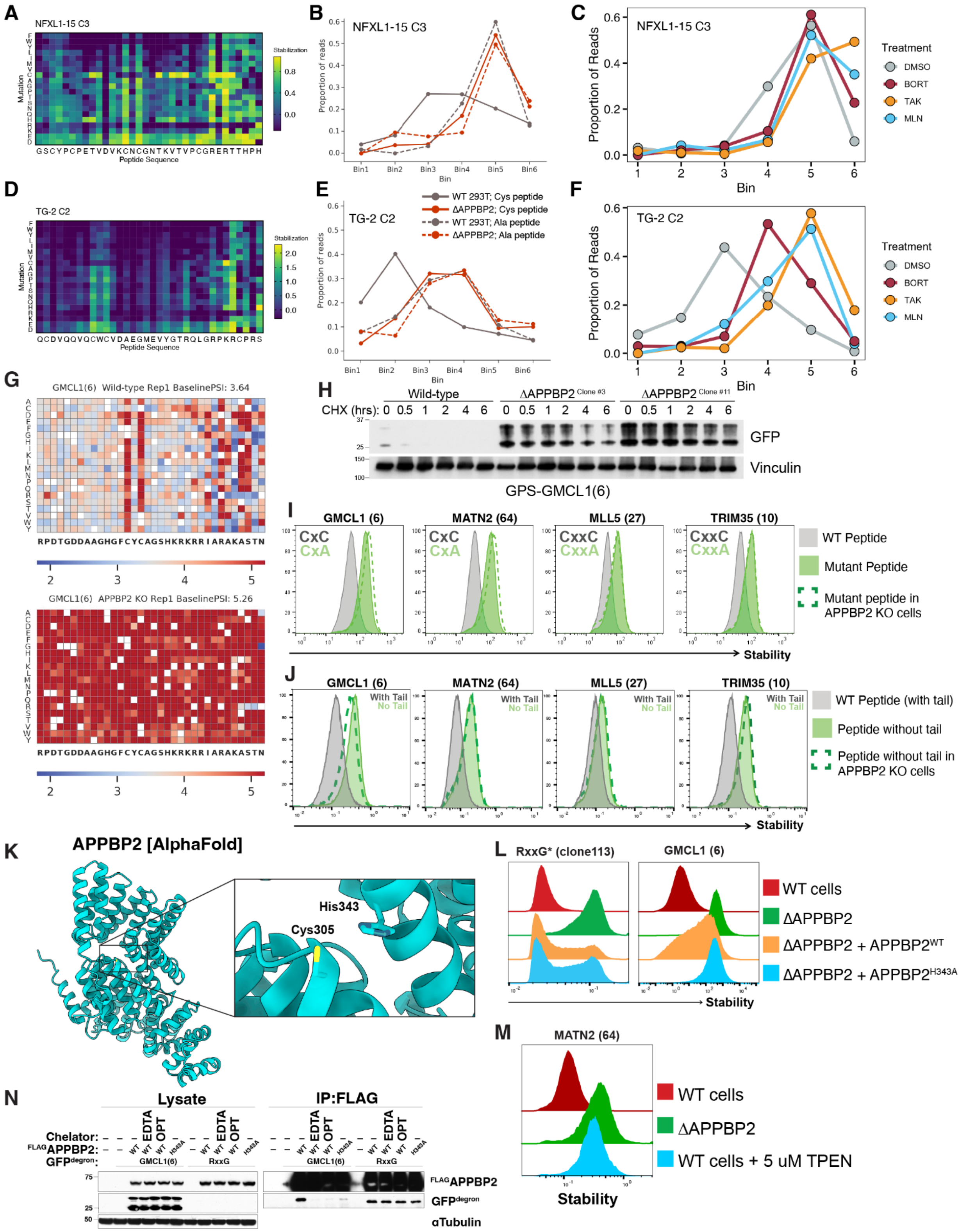

**Supplementary Figure 5**

**Fig. S5. APPBP2 targets a bipartite degron composed of a paired cysteine motif and C-terminal elements.** (**A**) Site-saturation mutagenesis heatmap of the NFXL1-15 degron, a substrate of APPBP2. (**B**) Bin-level read proportions from the cysteine stability screen in wild-type versus APPBP2 knockout 293T cells, highlighting APPBP2-dependent degradation. (**C**) Bin-level read proportions from cysteine stability screen performed with pharmacological inhibitors. (**D**-**F**) Equivalent data for TG-2, another APPBP2-dependent substrate. (**G**) Site-saturating mutagenesis heatmap for GMCL1(6) in wild-type cells and APPBP2 knockout cells. The APPBP2 substrate GMCL1(6) was reported in Timms et al., 2023 (*18*) (**H**) Cycloheximide chase assay of the GMCL1(6) degron in wild-type and APPBP2 knockout cells. The doublet is likely due to incomplete denaturation of GFP during SDS-PAGE. (**I**) Individual validation data for APPBP2 substrates that were reported in Timms et al., 2023 (*18*). Both cysteines in the motif appear to be required for APPBP2-dependent degradation. (**J**) Degradation of cysteine-containing peptides with or without the constant C-terminal tail (-RIARAKASTN*) used in Timms et al., 2023. The C-terminal sequence is critical to APPBP2 recognition. (**K**) AlphaFold 2 structure of APPBP2 showing His343 (critical for substrate interaction) and Cys305 (untested). (**L**) Stability assays of the GMCL1(6) degron in APPBP2 knockout cells reconstituted with wild-type or catalytic mutant (H343A) APPBP2. Clone113, a canonical RxxG degron substrate, serves as a control. (**M**) Stability of the MATN2(64) degron was assessed in APPBP2 knockout cells and upon treatment with TPEN. (**N**) Co-immunoprecipitation of GFP-tagged GMCL1(6) or clone113 with FLAG-APPBP2. Interactions were tested in the presence of the metal chelators EDTA or o-phenanthroline.

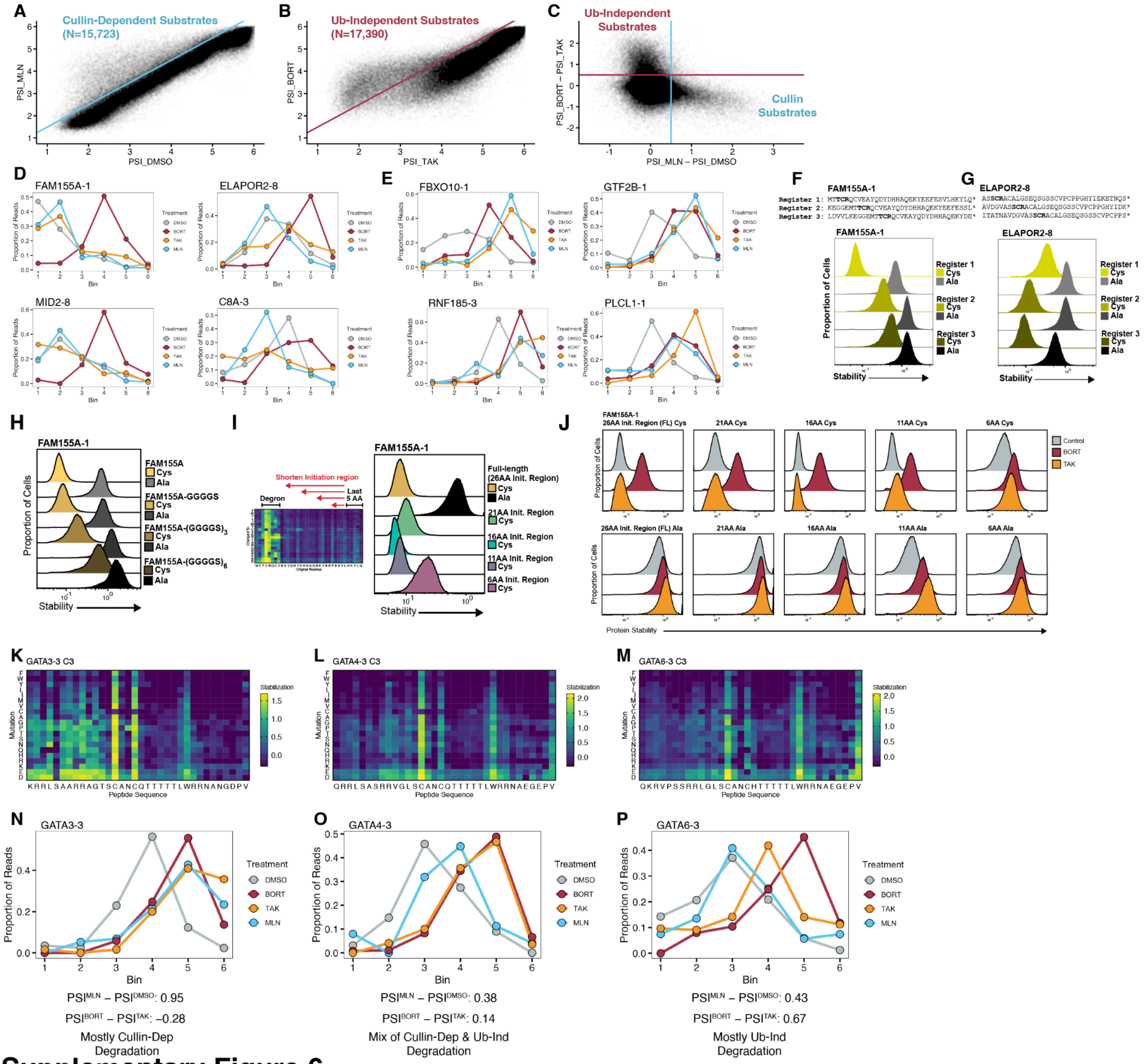


### Supplementary Figure 6

**Fig. S6. Sequence determinants of ubiquitin-independent degradation.** (**A**) Stability of all peptides under control conditions (PSI_DMSO) are compared to stability under Cullin inhibition (PSI_MLN). Using the threshold PSI_MLN - PSI_DMSO ≥ 0.5, 15,723 peptides are classified as Cullin-dependent. (**B**) Same analysis as in (A), comparing stability under bortezomib versus TAK-243. Using a cutoff of PSI_BORT − PSI_TAK ≥ 0.5, 17,390 peptides are classified as ubiquitin-independent substrates. (**C**) PSI_BORT − PSI_TAK is plotted against PSI_MLN − PSI_DMSO, revealing that Cullin-dependent and ubiquitin-independent substrates largely form non-overlapping groups. (**D**, **E**) Bin-level read proportions from the pharmacological screen, showing representative ubiquitin-independent substrates (D) and Cullin-dependent substrates (E). (**F**, **G**) GPS reporter assays of three overlapping tiles (registers) spanning FAM155A-1 (F) and ELAPOR2-8 (G). (**H**) C-terminal glycine-serine (GS) repeats appended to FAM155A-1 progressively stabilized the reporter in a length-dependent fashion, consistent with inhibition of proteasomal initiation. (**I, J**) Systematic truncation of the C-terminal tail of FAM155A-1 revealed that as few as 11 unstructured residues were sufficient to support degradation (I). Reporter assays were also conducted in the presence of bortezomib and TAK-243 (J), which revealed that shortening the tail to 6 residues abolished ubiquitin-independent degradation. (**K-M**) Site-saturation mutagenesis heatmaps of homologous peptides from GATA3 (K), GATA4 (L), and GATA6 (M). (**N-P**) Bin-level read distributions for GATA3-3, GATA4-3, and GATA6-3 under pharmacological inhibition. GATA3-3 is primarily degraded through the Cullin pathway, while GATA6-3 is predominantly processed via the ubiquitin-independent pathway.

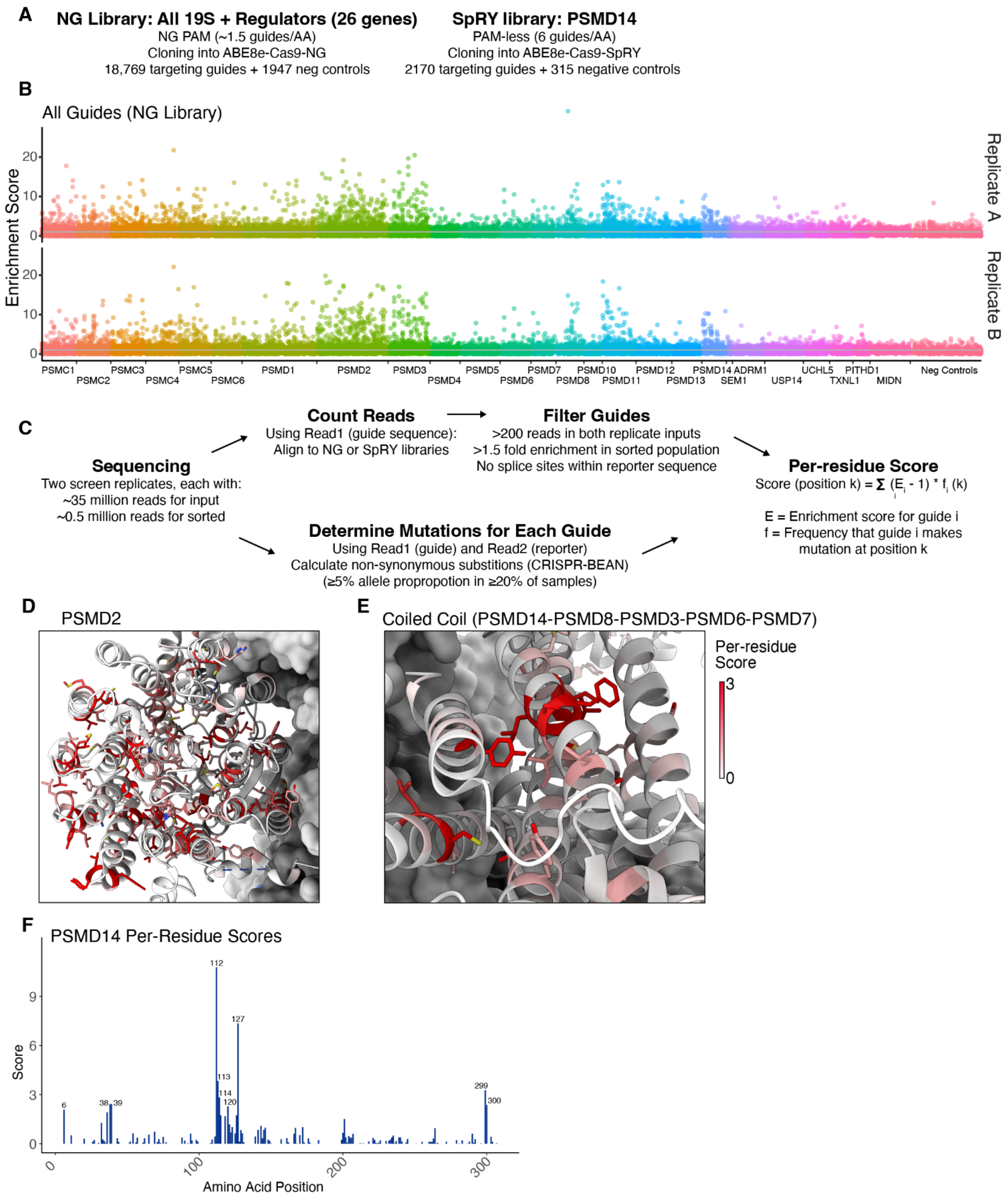


### Supplementary Figure 7

**Fig. S7. Design and additional results of base editor tiling screen.** (**A**) We used two base editor guide sub-libraries. The Cas9-NG library tiled across all 19S genes, while the Cas9-SpRY library targeted every nucleotide of PSMD14. (**B**) Overall enrichment scores for each guide in the Cas9-NG library. Two replicates were performed. Many of the top-scoring guides targeted splice sites, creating complete loss of function mutations. (**C**) By determining the mutations generated for each guide, we could calculate a per-residue score for each position in the 19S regulatory particle (RP). (**D**) High-scoring residues in PSMD2 clustered were typically buried or making hydrophobic contacts. Residues with scores of 0.5 and above are shown in stick representation. (**E**) High-scoring residues in the coiled coil region of the lid subcomplex within the 19S RP. (**F**) Per-residue scores for PSMD14. The active site, encompassing residues ~110-130, is the top-scoring region.

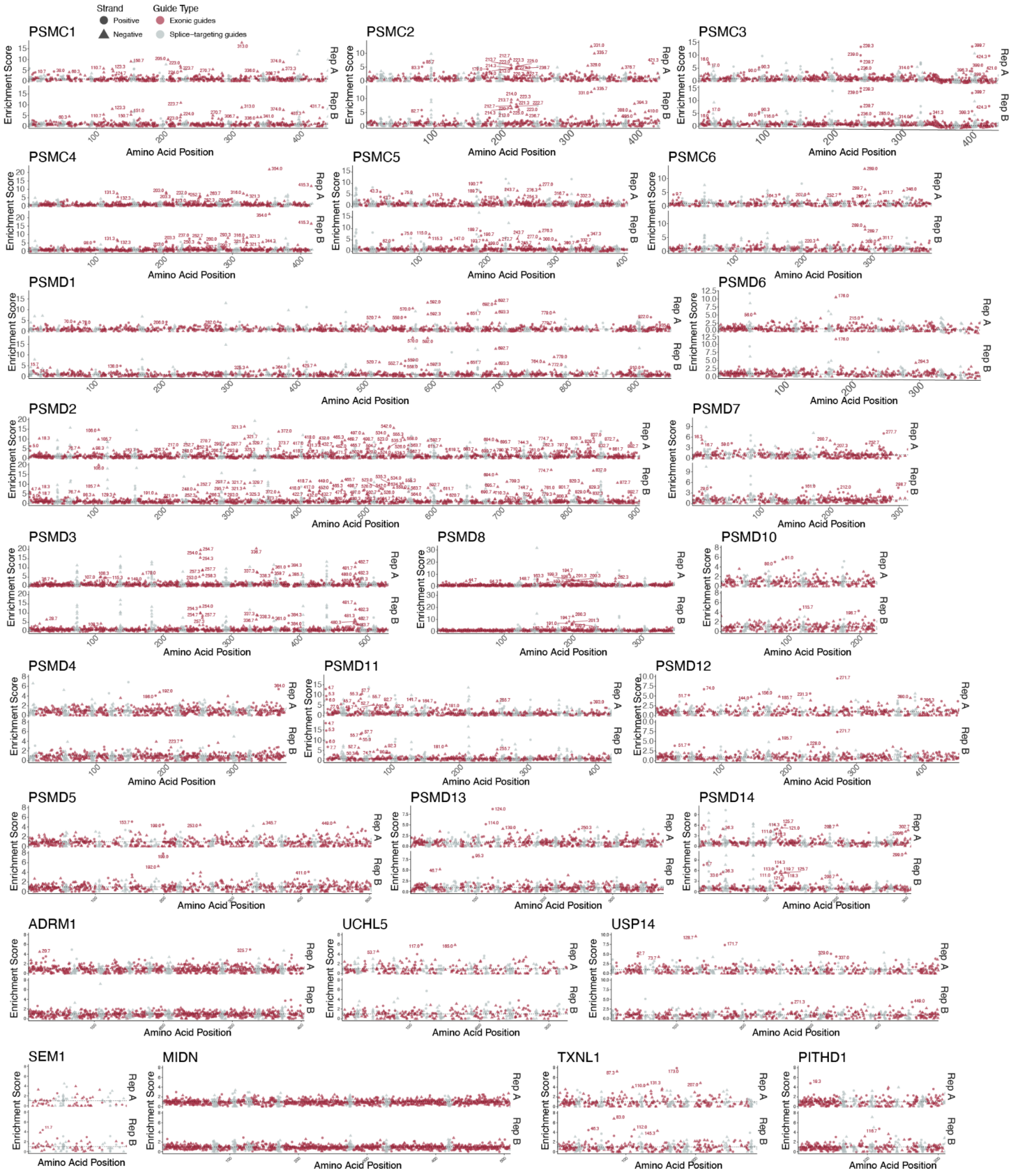


### Supplementary Figure 8

**Fig. S8. Guide-level results for base editor tiling screen.** Enrichment scores for each guide in the Cas9-NG library are shown, separated by target gene. Numbers refer to the amino acid that is contacted by position 6 of the protospacer. Positive strand-targeting guides are circular, while negative strand-targeting guides are triangular. Gray dots represent guides that target splice sites, whereas red dots target exonic regions.

**A**

| Peptide | Sequence | $IC_{50}$,Human PSMD14/PSMD7 | $IC_{50}$, Yeast RPN11/RPN8 |
|---|---|---|---|
| FAM Cys | MTT**C**RQCVEAYQDYDHHAQEKYEEFESVLHKYLQ | 7.1 µM | 4.8 µM |
| FAM Ala | MTT**A**RQCVEAYQDYDHHAQEKYEEFESVLHKYLQ | >100 µM | >100 µM |
| ELA Cys | AVDGVASS**C**RACALGSEQSGSSCVPCPPGHYIDK | 3.2 µM | 1.2 µM |
| ELA Ala | AVDGVASS**A**RACALGSEQSGSSCVPCPPGHYIDK | 13 µM | 90 µM |
| ELA NoCys | AVDGVASS**A**RA**A**ALGSEQSGSS**A**VP**A**PPGHYIDK | >200 µM | n.t. |
| ODC Cys | FPPEVEEQDASTLPVS**C**AWESGMKRHRAACASASINV | 72 µM | 60 µM |
| ODC Ala | FPPEVEEQDASTLPVS**A**AWESGMKRHRAACASASINV | >200 µM | >200 µM |

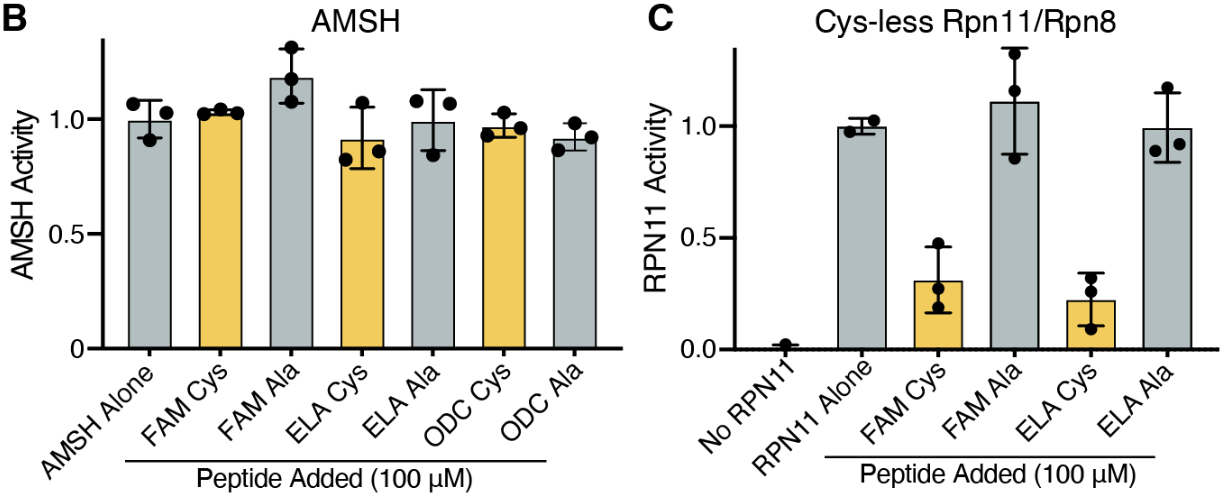


### Supplementary Figure 9

**Fig. S9. Cysteine-containing peptides specifically inhibit Rpn11.** (**A**) Half-maximal inhibitory concentrations ($IC_{50}$) of synthetic peptides against human PSMD14/PSMD7 and yeast Rpn11/Rpn8 dimers. Note that the "Ala" peptides contain mutations at the critical cysteine site but retain additional cysteines elsewhere in the sequence. n.t., not tested. (**B**) Cysteine-containing peptides do not inhibit AMSH, a JAMM-family deubiquitinase, even at concentrations of 100 µM. (**C**) Cysteine peptides still inhibit a cysteine-less Rpn11/Rpn8 dimer in which all cysteine residues were mutated to alanine, suggesting that inhibition does not occur via disulfide bonding to Rpn11 or Rpn8 cysteines.

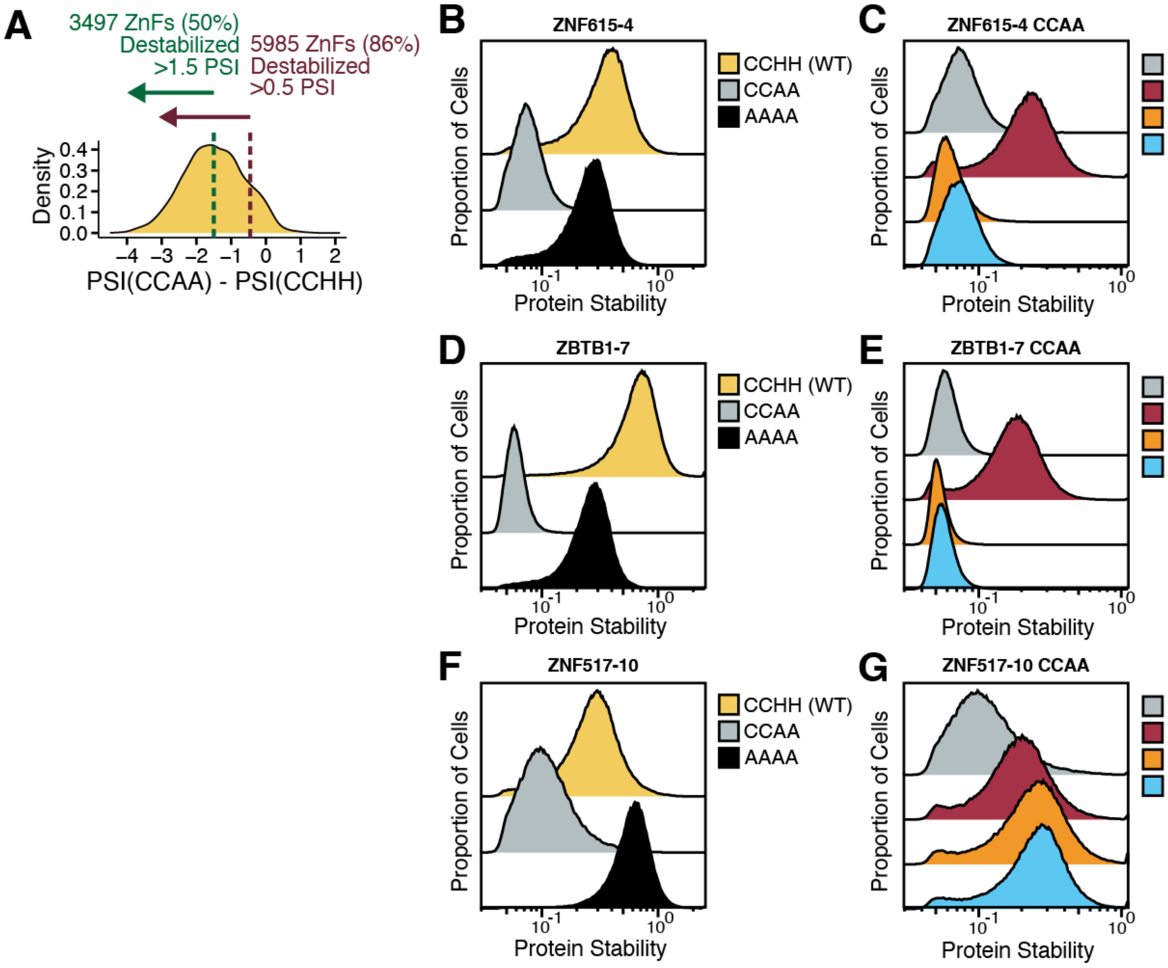


### Supplementary Figure 10

**Fig. S10. Additional zinc fingers that are degraded through cysteine-dependent pathways.** (**A**) Histogram showing the change in stability for ~7,000 zinc finger domains upon mutation of the two histidines used to fold around zinc. Wild-type CCHH domains were compared to CCAA mutants. (**B**-**G**) Individual stability reporter assays were used to determine the cysteine dependence (B, D, F) and degradation pathway (C, E, G) of selected zinc finger hits. ZNF615-4 (B, C) and ZBTB1-7 (D, E) were degraded via the ubiquitin-independent pathway, while ZNF517-10 (F, G) was Cullin-dependent.

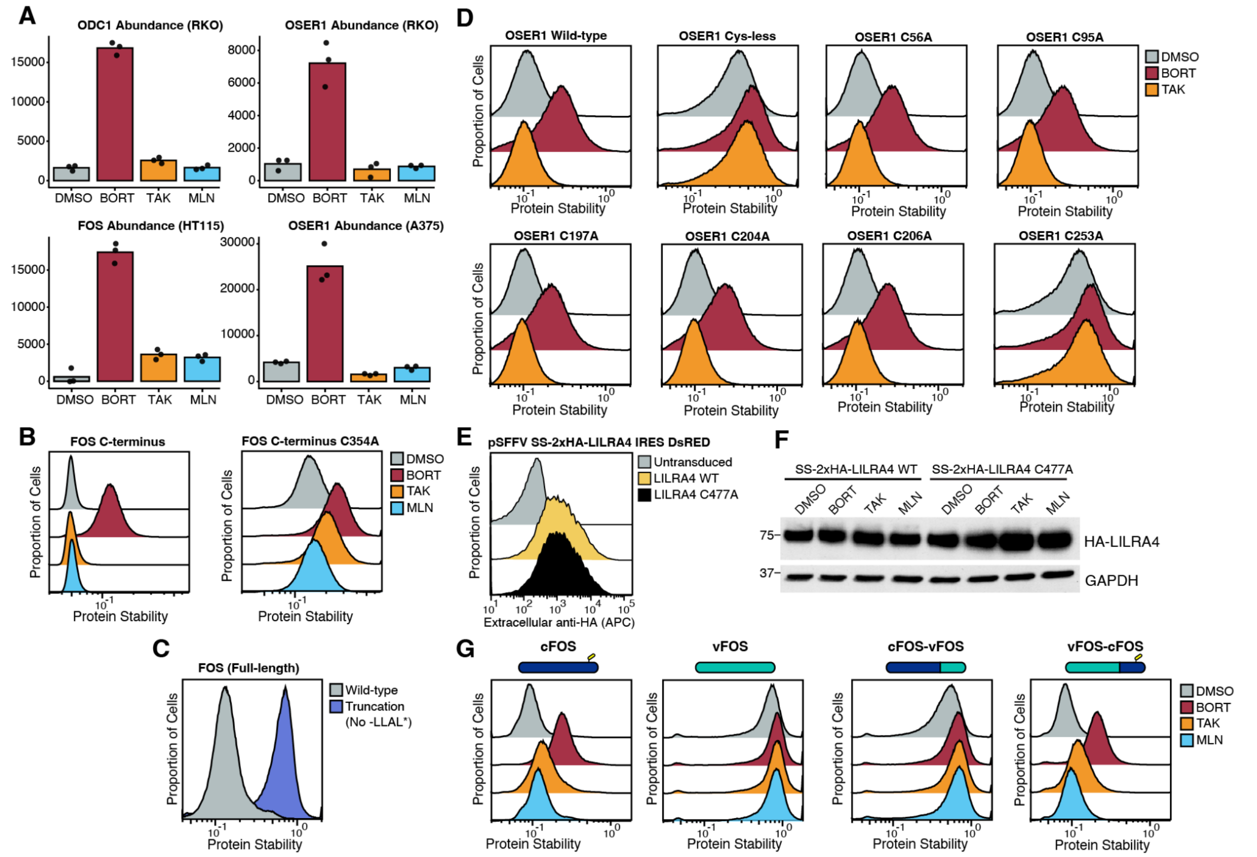


### Supplementary Figure 11

**Fig. S11. Quality control and regulatory substrates of the ubiquitin-independent pathway.** (**A**) Whole-cell proteomics in RKO, HT115, and A375 cell lines revealed that endogenous ODC1, OSER1, and FOS are stabilized by bortezomib treatment but not by TAK-243 or MLN-4924. Each point reflects a replicate. (**B**) The C-terminal 43 amino acids of FOS are sufficient to confer cysteine-dependent, ubiquitin-independent degradation, as shown using GPS reporter assays with the wild-type and cysteine-mutant (C354A) sequences. (**C**) Stability of full-length FOS and a truncation lacking the final four amino acids. (**D**) OSER1 mutants were assessed for stability in the presence of bortezomib or TAK-243. (**E**) Intact HEK-293T cells stably expressing 2xHA-tagged LILRA4 were assessed for plasma membrane localization using an anti-HA antibody. The 2xHA tag is after the signal sequence (SS), allowing for proper membrane localization. LILRA4 cells were gated on DsRED, which is expressed from an internal ribosome entry site (IRES) on the same construct. (**F**) The 2xHA-tagged LILRA4 is not appreciably degraded in HEK-293T cells, as assessed by western blotting. (**G**) Domain swapping experiments between cellular FOS (cFOS) and viral FOS (vFOS) indicates that the C-terminal tail (residues 333–380 of cFOS or 333–381 of vFOS) governs protein stability. This region harbors the ubiquitin-independent degron and may additionally confer ubiquitin-dependent degradation.

**Data S1. GPS stability screening of cysteine-containing peptides.**
Global Protein Stability (GPS) screening was performed in wild-type HEK293T and APPBP2 knockout cells, each in biological duplicate. The dataset includes Protein Stability Index (PSI) scores for wild-type ("c") and cysteine-to-alanine mutant ("a") versions of each peptide. The mutated cysteine appears at position 4, 9, or 14 depending on the peptide window (register 1, 2, or 3, respectively).

**Data S2. GPS site-saturating mutagenesis screening for 66 cysteine substrates.**
This dataset includes bin counts and PSI scores for every single-substitution variant of 66 cysteine degrons.

**Data S3. CRISPR knockout screening of 16 cysteine substrates using ubiquitin-wide guide library.**
Gene- and guide-level enrichment results are provided from MAGeCK analysis of 16 different CRISPR knockout screens using the ubiquitin-related CRISPR library. The full library of sgRNA sequences is also included.

**Data S4. CRISPR knockout screening of 3 cysteine substrates using genome-wide guide library.**
Gene- and guide-level enrichment results from MAGeCK analysis are provided for genome-wide CRISPR knockout screens using the Root library. Guide sequences are also included.

**Data S5. GPS stability screening of cysteine-containing peptides in the presence of ubiquitin-proteasome inhibitors.**
PSI scores and normalized bin counts are provided for GPS screens conducted in the presence of DMSO, bortezomib, MLN4924, and TAK-243.

**Data S6. CRISPR base editor screening of the 19S regulatory particle.**
This dataset includes mutational outcomes, read counts, and per-residue functional scores from tiling base editor screens (NG and SpRY libraries) targeting genes encoding 19S proteasome subunits.

**Data S7. GPS stability screening of ~7,000 zinc finger domains.**
PSI scores for wild-type (CCHH) and mutant (CCAA, HisAla) zinc finger peptides, in cells treated with DMSO, bortezomib, TAK-243, or MLN-4924.

**Data S8. Whole-cell proteomics in the presence of ubiquitin-proteasome inhibitors.**
Mass spectrometry-based proteomics data measuring total protein abundance in RKO, HT115, and A375 cells treated with DMSO, bortezomib, TAK-243, or MLN-4924.
